# Inferring scaling laws of branching networks from their geometry by global optimization

**DOI:** 10.64898/2026.10.30.763120

**Authors:** Etienne Jessen, Marc C. Steinbach, Dominik Schillinger

## Abstract

Optimality principles are widely used to understand branching networks, yet they are tested on individual branchings that cannot reliably distinguish competing scaling laws. We formulate the missing link between the laws and measurements by mapping each law to its predicted branch points through global optimization. Vascular branch points closely match weighted-length optima when each vessel is allowed its own fitted weight. Inferred scaling laws reveal type-specific geometric signatures hidden from local analysis. They beat all tested laws, including Murray’s law in 1,679 of 1,744 held-out hepatic and pulmonary trees. For coronary arteries on the heart surface, an exact discrete optimization model extends the framework beyond convex geometry and supports area-preserving scaling. The framework also identifies distinct laws for airways and plant shoots while rejecting weighted-length optimality for neuronal axons. Global optimization turns network geometry into a quantitative test of the principles that govern branching networks.

---

For a century, optimality principles have been used to explain the structure of biological networks such as the cardiovascular system. In 1926, Murray linked vascular design to transport efficiency and described the optimal angle [1] and radii [2] at a branching. Murray’s law motivated many studies on morphology and topology in biological trees, ranging from blood vessels and airways [3–7] to river basins [8], neuronal arbors [9, 10] and plant shoots [11].

Most optimality principles for biological trees are expressed as weighted-length problems. A scaling law then sets the weights across a tree’s hierarchy. The weights correspond to the physiological effects researchers deem most important. In Murray’s law they are the vessels’ cross-sections, derived from minimizing the cost of pumping and maintaining blood. Other laws followed, for example by adding the effect of turbulent flow [12], minimizing wave reflections in large arteries [3, 13], or resisting wind-induced stresses in tree branches [14]. However, competing laws now exist for one system, as in the coronary arteries, where the law of Huo and Kassab [15] rivals Murray’s law and area preservation [16].

Whether any proposed law holds is still debated. A meta-analysis in coronary trees [16] showed high heterogeneity between studies. Recently, Meng et al. found that the branchings of six biological networks depart systematically from the predictions of (unweighted) length and volume minimization [17]. However, these validations mostly rely on local analysis of radii and angles at isolated branchings and are pooled over common hierarchical features like Strahler order [16–19]. Yet networks of different form can share similar statistics [8], and local tests cannot tell different laws apart. This problem is made worse because the scaling parameter derived from the radii of similar vessels varies widely between studies [15, 16, 18, 19].

Here, we propose a new computational framework to test and infer scaling laws from biological trees. Instead of relying on local analysis and radii, we use global optimization on the centerlines of whole trees. We do not derive weights from assumptions but ask which weights fit measured centerlines best. We hypothesize that the topology and centerlines of large- and intermediate-scale vessels form a skeleton that changes more slowly than the radii. In contrast to Meng et al. [17], we further hypothesize that the weighted length describes the skeleton across organs and vessel types, which differ only in their scaling law.

Testing a law on a whole tree requires predicting the optimal geometry (positions of all branch points) under that law. We achieve this by casting this prediction into a convex optimization problem [20], which has exactly one optimal geometry per law and is solved in milliseconds even for large trees. This allows us to infer scaling laws directly from the centerlines by solving an inverse optimization problem [21]. On the curved heart surface, we solve the prediction exactly by dynamic programming over the tree [22], with one shortest-path search over the surface mesh per vessel. To our knowledge, we are thus the first to infer a vascular scaling law from the full geometry of whole trees.

We find that the geometries of real vascular trees [23–28] lie near the optimum of a weighted length. Furthermore, we can infer scaling laws that hold on held-out trees. Murray’s law fails in four of five vessel types, and the inferred law on the coronary arteries [29, 30] lies close to area preservation. The inferred laws can also tell the two trees of one organ apart. Finally, our framework infers laws for airways [26, 31] and plant shoots [11, 17], and rejects the weighted-length hypothesis for neuron axons [32, 33]. Our framework thus turns the geometry of measured trees into evidence for or against a design principle, in any network with a measured topology.

## Results

### Local analysis leaves scaling unresolved

We first test a scaling law as most studies do, by local analysis at isolated branchings. In vascular trees, Murray’s law [2] assumes that each vessel minimizes its cost of pumping and maintaining blood. At each branching it describes the optimal distribution of radii with radius exponent *γ* = 3 as

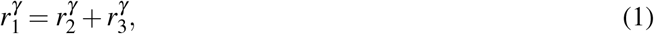

where *r*_1_ is the parent radius and *r*_2_ and *r*_3_ are the daughter radii. The same derivation predicts the optimal branching angles 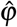 from the radii (Eq. (19)) [1].

To test these predictions, we measure local radius exponents *γ* and branching angles *ϕ* at the branchings of a pulmonary vein and a portal vein tree, grouped by Strahler order (Fig. 1a,b). Measurements scatter widely within each Strahler order (Fig. 1c,d). The optimal radius exponent is often based on the assumed flow regime, ranging from *γ* = 2 for pulsatile flow [3] through *γ* = 7*/*3 for turbulent flow [12] to *γ* = 3 [2] for laminar flow. Yet most radius exponents of the example trees lie outside this range (Fig. 1d), which no flow regime explains. Measured angles *ϕ* also correlate little with Murray’s predicted angles 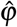 (Fig. 1e). The scatter persists across 2,559 trees of five vessel types (Methods), where the interquartile range of *γ* within one Strahler order of a tree spans 1.2 to 1.6. Taken together, local analysis of radii can neither settle whether a proposed scaling law holds nor infer one.

**Figure 1.**
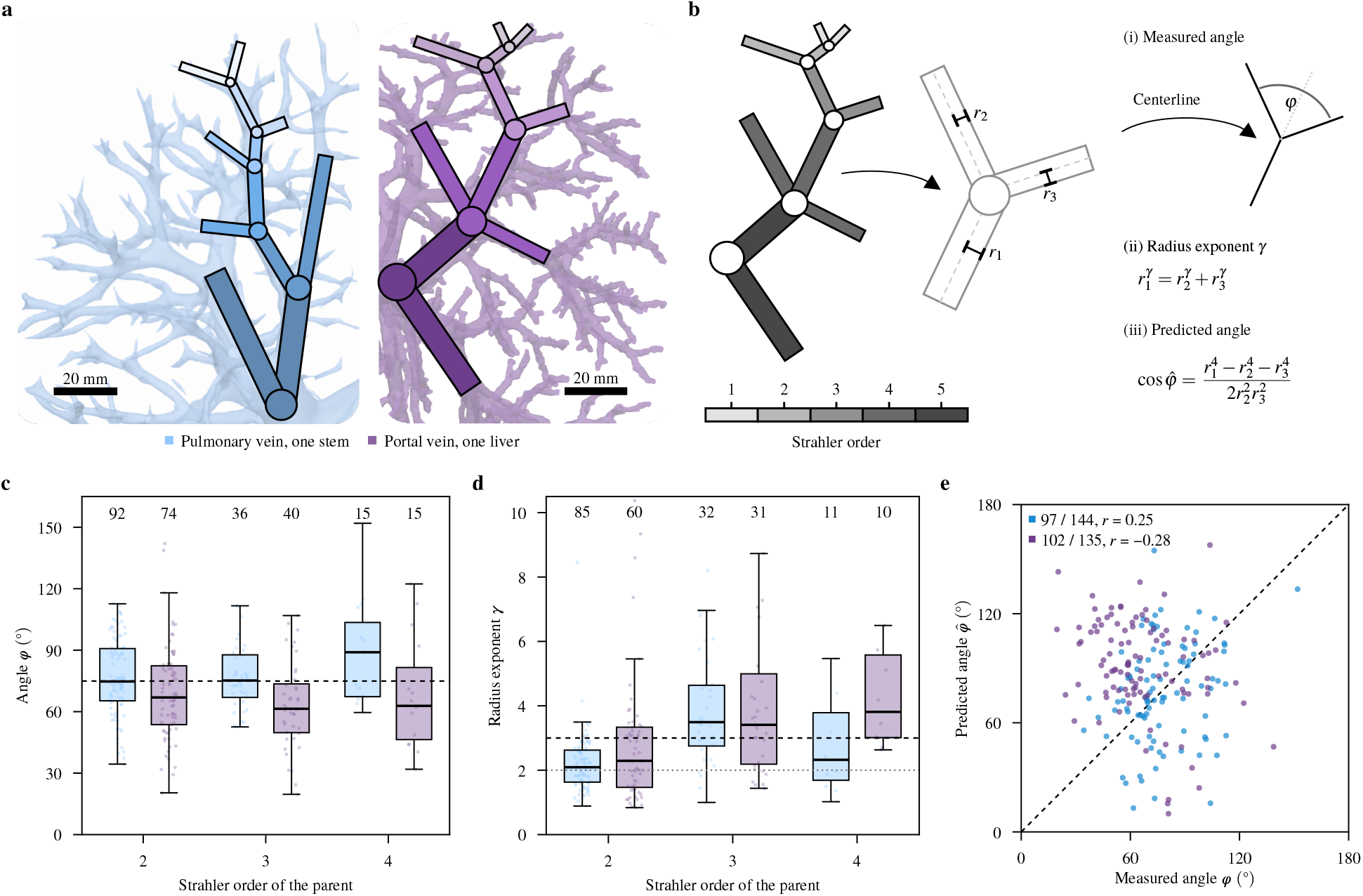
Local branching measurements versus Murray’s law. **a**, Pulmonary vein tree from one lung and portal vein tree from one liver, with schematic overlays of representative root-to-terminal routes. Scale bars, 20 mm. **b**, Branching measurements: daughter angle *ϕ*, radius exponent *γ* relating parent and daughter radii, and angle 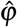 predicted by Murray’s law. **c**,**d**, Branchings where both daughters are thinner than the parent, grouped by parent Strahler order. Points represent branchings; numbers indicate sample sizes. Groups with fewer than eight observations are omitted. Boxes show quartiles, center lines medians, and whiskers the 1.5 interquartile ranges of the box limits. **c**, Measured angles. The dashed line marks Murray’s prediction of approximately 75^°^ for symmetric branching. **d**, Measured radius exponents. The dashed line marks Murray’s prediction *γ* = 3, and the dotted line *γ* = 2. Below it 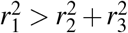, and the formula in **b** has no real angle. **e**, Predicted versus measured angles; the dashed line indicates equality. Annotations give the branchings with a real predicted angle, of all branchings, and Pearson’s *r*.

### Inferring scaling laws from global network geometry

A scaling law shapes not only a local branching but the global geometry of the entire network. We therefore read this geometry from the measured centerlines instead of the radii, which change with the hemodynamic state [34–36] and are measured to only about ± 10% accuracy [19]. In contrast, the centerlines offer a stable measurement target, which we expect to change more slowly (Fig. 2a).

**Figure 2.**
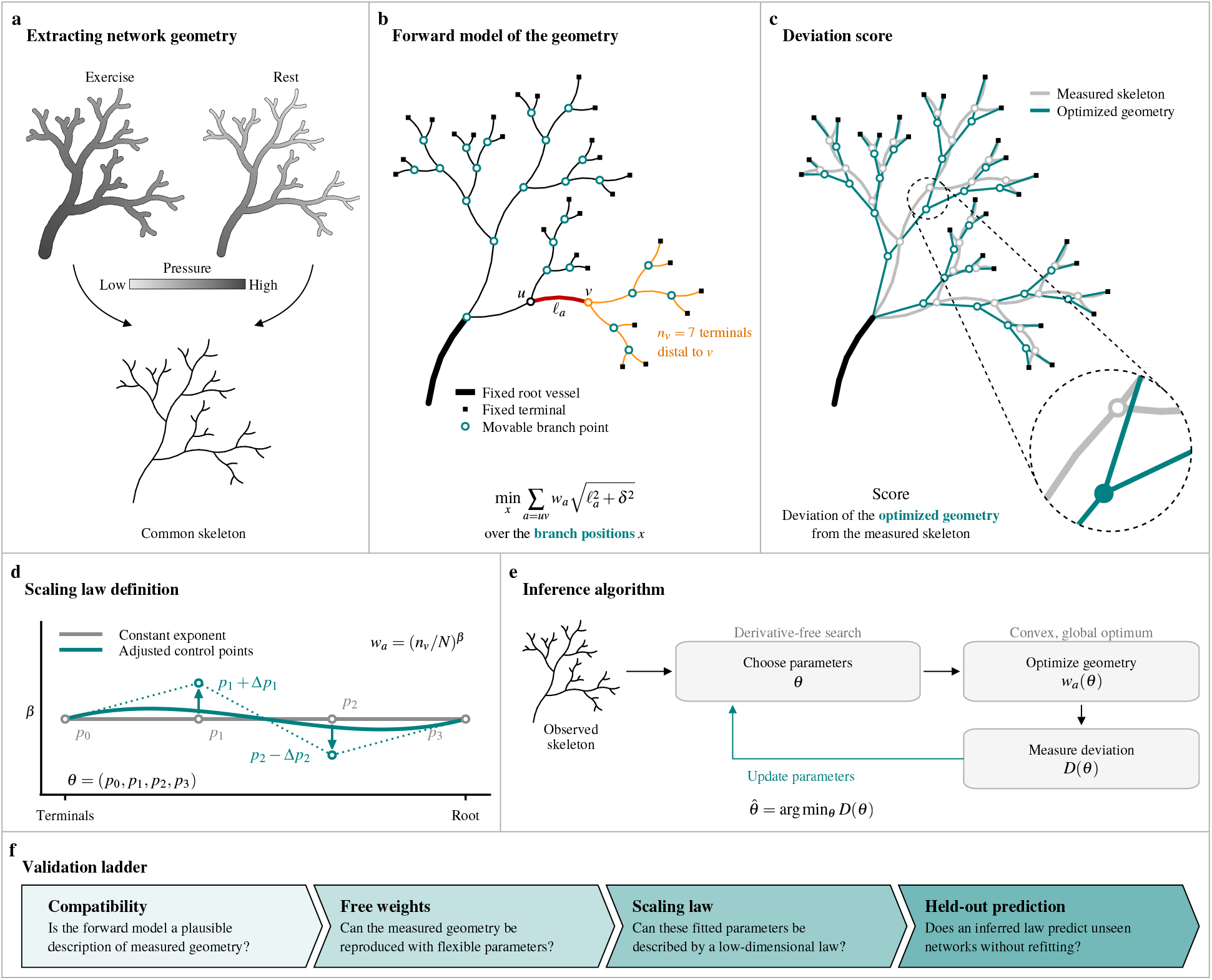
Inferring and testing scaling laws from tree geometry. **a**, A synthetic tree in two hemodynamic states, with vessel radii and pressure varying on a common skeleton. **b**, A forward model that describes the geometry with a weighted length problem over fixed topology, root vessel and terminal positions. Branch points are optimized under given weights, and *l*_*a*_ denotes vessel length. **c**, Measured skeleton and optimized geometry under Murray’s law. The inset illustrates branch-point displacement. Geometric deviation from the measured skeleton provides the score *D*. **d**, One candidate scaling law: *w*_*a*_ = (*n*_*v*_*/N*)^*β*^ for the vessel *a* ending at node *v*, where *n*_*v*_ counts the terminals distal to *v* and *N* all terminals. *β* can change across the hierarchy using a Bernstein curve with control points *θ* that can be fitted. **e**, Inference algorithm to identify the parameter *θ* yielding the smallest score *D*. **f**, Validation ladder with progressively stronger tests for a candidate law to describe given measurements. A failure on one step fails the entire ladder.

A network should lie near the optimal geometry of its scaling law. Our key methodological advance is to compare the measured geometry with the global optimum of a candidate law. A forward model tests a candidate by mapping it to one optimal geometry (Fig. 2c). It predicts every branch point of a whole tree at once, each compared directly with its measured position. To make this tractable, we keep the topology, root and terminals as measured. By using inverse optimization, this method allows us to infer the law from the geometry, whose predicted vessels lie closest to the measured ones (Fig. 2e).

For vascular trees, Murray’s law can be formulated as a weighted length (Fig. 2b) [2], where each weight is the vessel’s cross-sectional area. This weighted-length form is the basis of many other scaling laws in network modeling [9, 11]. Smoothing each vessel length by *δ* = 5 mm keeps short vessels from collapsing to a point (Supplementary Fig. 1) and makes the problem strictly convex. With equal flow per terminal, a scaling law sets each vessel’s weight from the share *n*_*v*_*/N* of terminals it supplies, *w*_*uv*_ = (*n*_*v*_*/N*)^*β*^ with *β* = 2*/γ*. Here *β* = 0 gives minimal length, *β* = 2*/*3 Murray’s law and *β* = 1 area preservation. Since West et al. predicted area preservation in large vessels and Murray’s law in small ones [3], we allow *β* to vary across a tree’s hierarchy (Fig. 2d). Strict convexity gives each law one global optimum, found in milliseconds. The inference framework was tested on synthetic trees under noise, where it recovered all prescribed laws (Supplementary Fig. 2).

A lower deviation score shows only that one law fits better than another, not whether the scaling law applies to a given network at all. Therefore, we propose a validation ladder of four tests of rising difficulty that any scaling law and forward model must pass (Fig. 2f). It starts with a compatibility check that tests whether the law holds locally at isolated branchings. Free weights, one per vessel, then ask whether any weighting places the whole network near its measured geometry. Only then should a proposed scaling law be fitted, and it must be judged on held-out networks. A law that fails one test fails the ladder.

### Vessel trees lie near weighted-length optima

The first test of the validation ladder asks whether the centerlines of human vascular trees are compatible with a weighted-length optimum (Fig. 2f). At a weighted-length optimum, each branch point lies inside the convex hull of its neighbors, since otherwise no positive weights balance the pull of its vessels. Since a measured branch point never lies exactly in the plane of its three neighbors, we call it *compatible* when it lies close to their triangle (Methods and Supplementary Fig. 3a). In livers and lungs, a median of 84.2% to 92.4% of branch points were compatible, against only around 20% for vessels placed in random directions (Fig. 3a).

**Figure 3.**
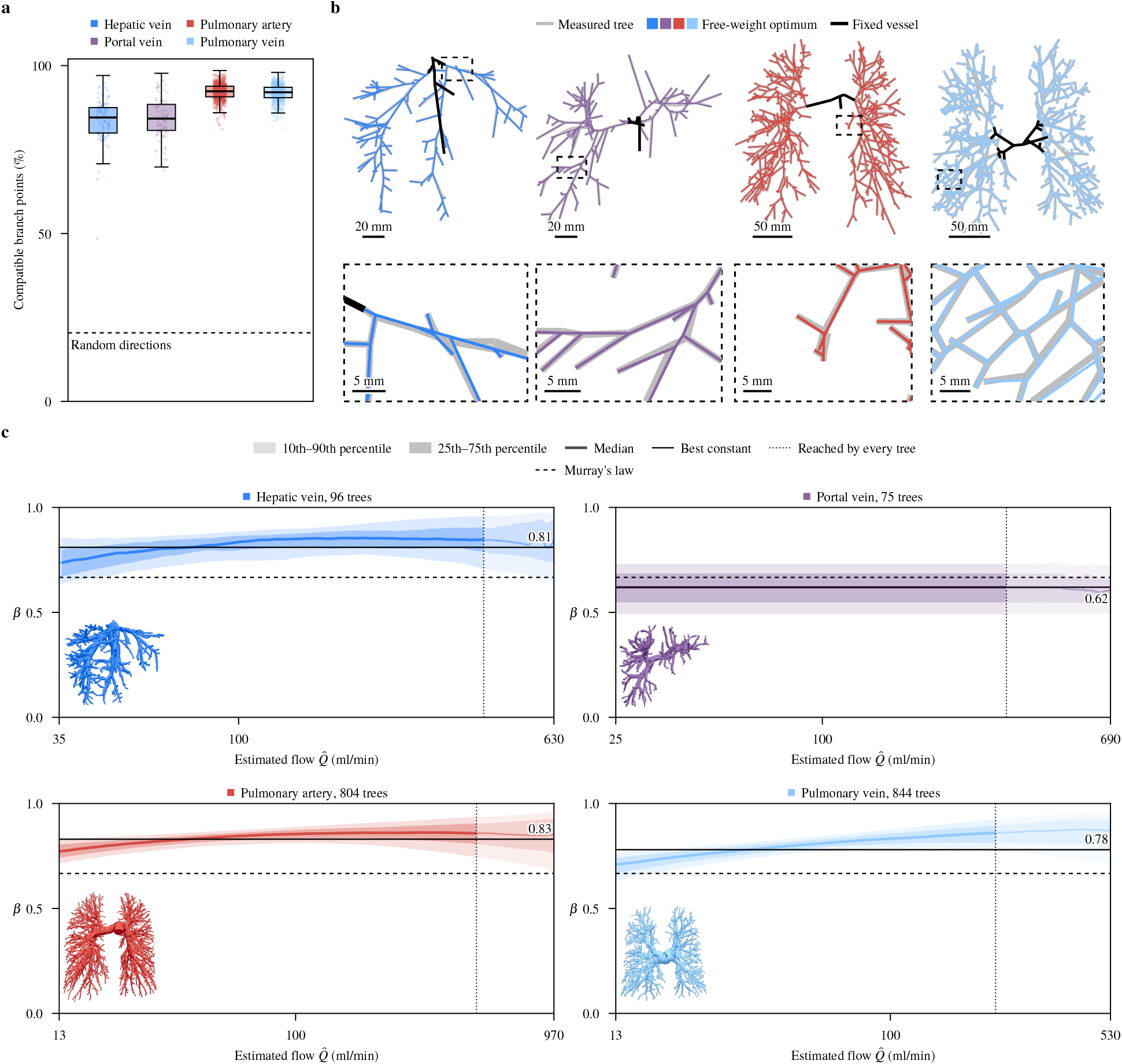
Vessel trees lie near weighted-length optima, and their laws depart from Murray’s. **a**, Share of each tree’s branch points that are compatible, per vessel type (*n* = 96 hepatic veins, 75 portal veins, 804 pulmonary arteries and 844 pulmonary veins); random directions keep the observed vessel lengths, pooled over the types. **b**, One measured tree per type under its free-weight optimum; insets zoom on the branch point of median displacement. **c**, Fitted laws of the four vessel types against the estimated flow 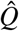, the organ’s root flow distributed by perfusion territory. Median and percentiles are taken across the trees of **a**.

Compatibility can rule out a weighted-length hypothesis for a network, but cannot say how far the network lies from the optimum. The second test therefore asks whether any positive weights make a whole measured tree a weighted-length optimum, a question open since Murray proposed that vessels minimize a cost [1]. Fitted to each tree (Methods), free weights place the branch points of the median tree of every vessel type about a tenth of a vessel length from the measured ones (deviation score 0.11 to 0.15, in sample; Fig. 3b). Copies of the same trees with random vessel directions lie about half a vessel length away. Vessel geometry is therefore far from random, and to our knowledge this is the first evidence that whole vascular trees lie near the optimum of a weighted length. Such optima are also plausibly reachable in biology, since in models of flow networks, local adaptation during growth finds near-global optima [37].

### Murray’s law fails in three vessel types

Fitting free weights only shows that weights exist for a tree, but one weight per vessel defines no law. The third test of the validation ladder asks whether a low-dimensional scaling law can also describe the measured geometry. Run on the pulmonary arteries and veins of the lung and the portal and hepatic veins of the liver, our inverse optimization gives each tree one law, whereas the local radius exponents of the same tree scatter widely (Fig. 1d). Of the commonly assumed weightings, Murray’s law fits best (Supplementary Fig. 4a). However, in hepatic veins, pulmonary arteries and pulmonary veins, the best constant *β* lies above it, at 0.78 to 0.83 (Fig. 3c; Supplementary Fig. 5b), with 95% bootstrap intervals over trees no wider than 0.02. Under Murray’s volume cost, these laws correspond to radius exponents 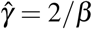 of 2.4 to 2.6, between area preservation (*γ* = 2) and Murray’s law (*γ* = 3). In contrast, portal veins stay close to Murray’s law, at *β* = 0.62. The fitted laws also tell the two trees of one organ apart, giving the pulmonary artery the larger *β* in 658 of 690 lungs and the hepatic vein in 57 of 59 livers.

**Figure 4.**
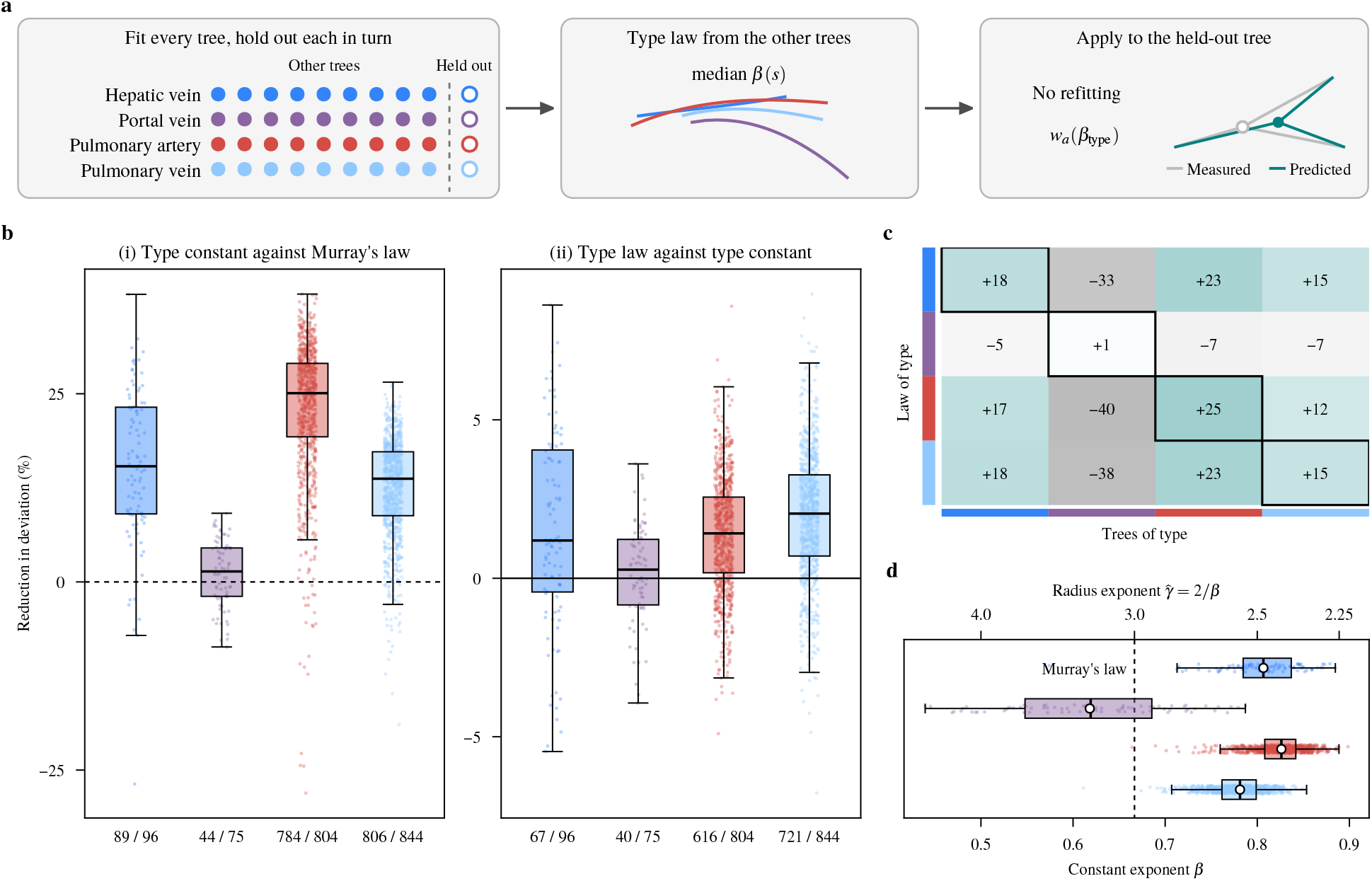
Held-out test and transfer of type laws. **a**, Held-out test. Each tree in turn is held out, and its type law *β*_type_, the median at each *s* of the fitted laws of all other trees of its type, predicts its geometry without refitting. **b**, Reduction in deviation per held-out tree for hepatic veins (*n* = 96 trees), portal veins (*n* = 75), pulmonary arteries (*n* = 804) and pulmonary veins (*n* = 844). Fractions denote how many trees improved. Boxes show quartiles and medians, and whiskers reach at most 1.5 interquartile ranges beyond the box. **c**, Reduction in mean deviation against Murray’s law when the type law of each row is applied to the trees of each column (teal better, gray worse). **d**, Constant exponent *β* of each tree (boxes as in **b**) and the type constant (open circles).

**Figure 5.**
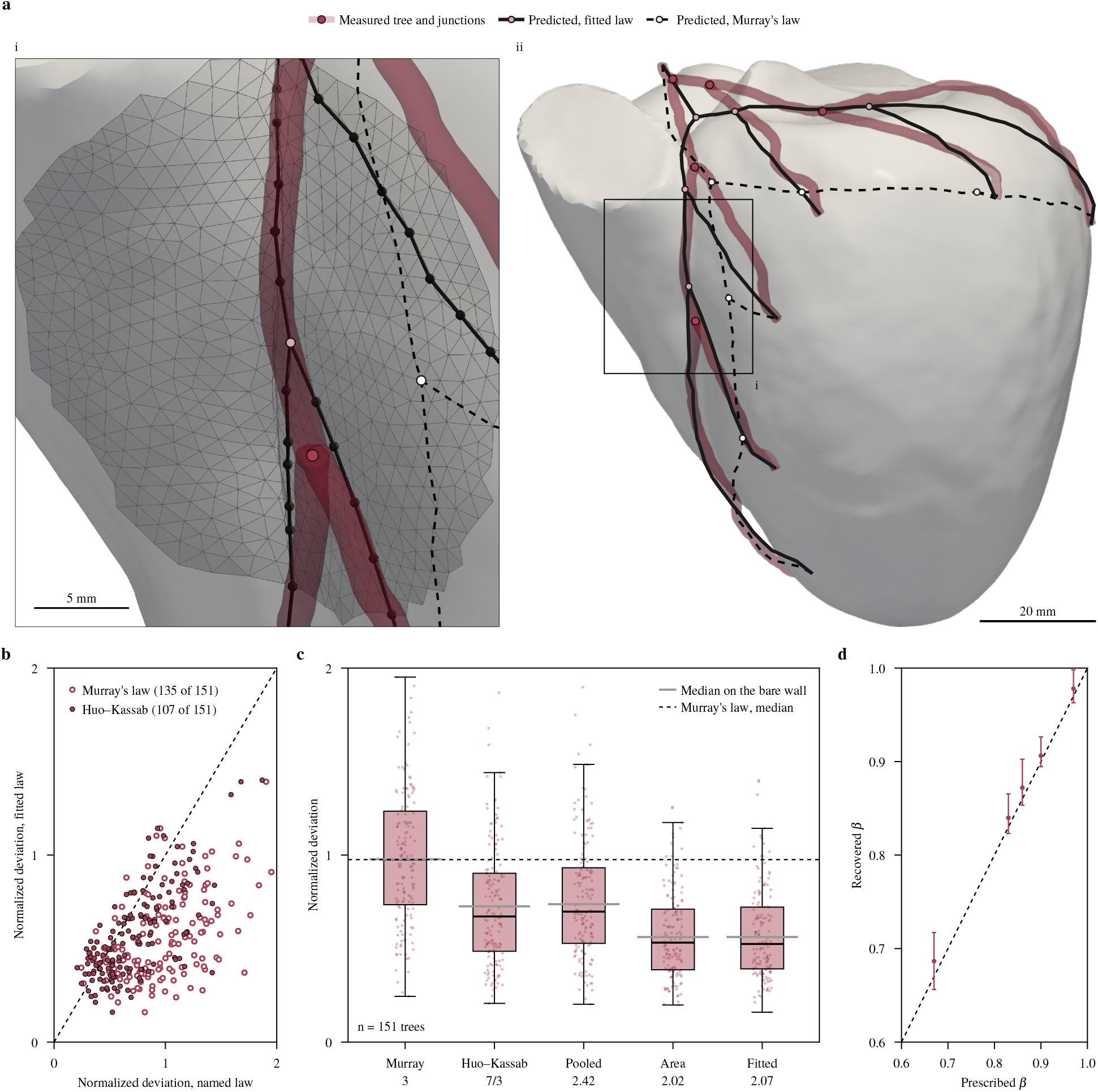
Coronary trees on the heart surface. **a**, (i) The window marked in (ii), magnified. Predicted vessels run along shortest paths over a surface 2.5 mm off the heart wall (gray triangles). (ii) A held-out left coronary tree, measured and predicted under the fitted law and Murray’s law. Roots lie a median of 15 mm above the wall. **b**, Normalized deviation per held-out tree under the fitted law (*β* = 0.97) against Murray’s law and Huo and Kassab’s law [15]. The dashed line marks equality. **c**, Normalized deviation under five laws, labeled by the radius exponent 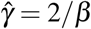 (pooled, the coronary exponent pooled over studies [16]). Boxes show quartiles and medians, and whiskers reach at most 1.5 interquartile ranges beyond the box. **d**, Recovered against prescribed *β* on synthetic trees (median and 10th–90th percentile over 50 trees). Deviations are normalized by the median vessel length (25.87 mm); *n* = 151 held-out trees.

A law that varies along the tree needs more parameters. To prevent overfitting, each type takes only as many control points *M* as it needs to lower the deviation on its held-out trees (Methods). Pulmonary trees support three, hepatic veins four and portal veins one, so the portal geometry shows no change of *β* along the tree (Supplementary Table 1). The varying laws show one shared signature in pulmonary arteries, pulmonary veins and hepatic veins (Fig. 3c). Their median *β* rises from 0.70–0.77 at the small vessels to 0.84–0.85 near the root. Noise alone rarely changes a law this much (Supplementary Fig. 2). Exponents that change with vessel size were measured from radii [38] and predicted from wave reflection [13], but our laws are directly inferred from the positions of branch points.

### Scaling laws transfer between organs

The last test of the validation ladder asks whether a type’s law predicts trees it was never fitted to (Fig. 4a). The *type constant* of a tree, the median constant *β* of all other trees of its type (Fig. 4d), beats Murray’s law in 1,679 of the 1,744 held-out trees of the three types, by 13 to 23% (Fig. 4b(i)). In portal veins, it beats Murray’s law in only 44 of 75 trees (sign test, *p* = 0.17). Letting *β* change along the tree lowers the deviation by a further 1.4 to 2.0% per tree in the three types, but not in portal veins (Fig. 4b(ii)).

A stronger test asks whether a law fitted in one organ predicts the trees of another, scanned in other datasets (Supplementary Table 2). The law of the hepatic veins predicts the lung trees almost as well as their own type laws (Fig. 4c). On pulmonary arteries it takes 23.1% off the deviation under Murray’s law, against 25.1% for their own law, and on pulmonary veins it equals their own. In return, the laws of both lung trees predict hepatic veins about as well as the hepatic veins’ own law. Most of what transfers is the mean of *β*, and its rise along the tree adds 2.4 to 2.7 percentage points between hepatic and pulmonary veins. The portal-vein law predicts every other type worse than Murray’s law, and is predicted worse by any other type (Fig. 4c).

The transferred laws also beat the closest law from the literature that we tested, the *γ* = 7*/*3 of Huo and Kassab [15] (*β* = 0.86 under a volume cost), which takes only 2.1% off on pulmonary veins. Huo and Kassab refitted their exponent in each tree, whereas our laws predict another organ’s trees without refitting.

### Coronary geometry favors area preservation

In coronary arteries, many rival laws exist. Murray’s law, Huo and Kassab’s *γ* = 7*/*3 [15] and area preservation are all used. Radii cannot decide between them, since radius exponents scatter across studies from 1.1 to 3.53, a spread partly ascribed to imaging resolution [16].

Since coronary arteries run over the heart wall, we need to replace the convex model with a discrete one, which keeps each free branch point on a surface off the wall and runs each vessel along its shortest path over it (Fig. 5a(i)). Dynamic programming finds its exact optimum over the surface mesh (Methods), and the fit and the held-out test carry over. As the fit recovers prescribed laws on synthetic trees (Fig. 5d), we fit the law on 151 left coronary trees from ImageCAS-X [29, 30], holding each out in turn.

The geometry decides between these laws. The fitted law, *β* = 0.97, halves the median deviation of Murray’s law and beats it in 135 of the 151 held-out trees (Fig. 5a(ii)–c). It also beats Huo and Kassab’s law in 107 of them (Fig. 5b). Only area preservation (*β* = 1) fits as well, so the geometry rejects Murray’s and Huo and Kassab’s laws. The fit without smoothing gives *β* = 0.91, so 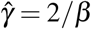 is at most about 2.2, below the *γ* = 2.42 measured in human coronary arteries [16]. To our knowledge, this is the first coronary law inferred from whole three-dimensional trees on the heart surface and judged on held-out trees, whereas an earlier study fitted small planar trees, in sample [39].

### Beyond blood vessels

As the framework works with any branching network with measured centerlines and topology, we run it on five networks beyond blood vessels (Fig. 6a). Each network goes through the validation ladder (Fig. 2f).

**Figure 6.**
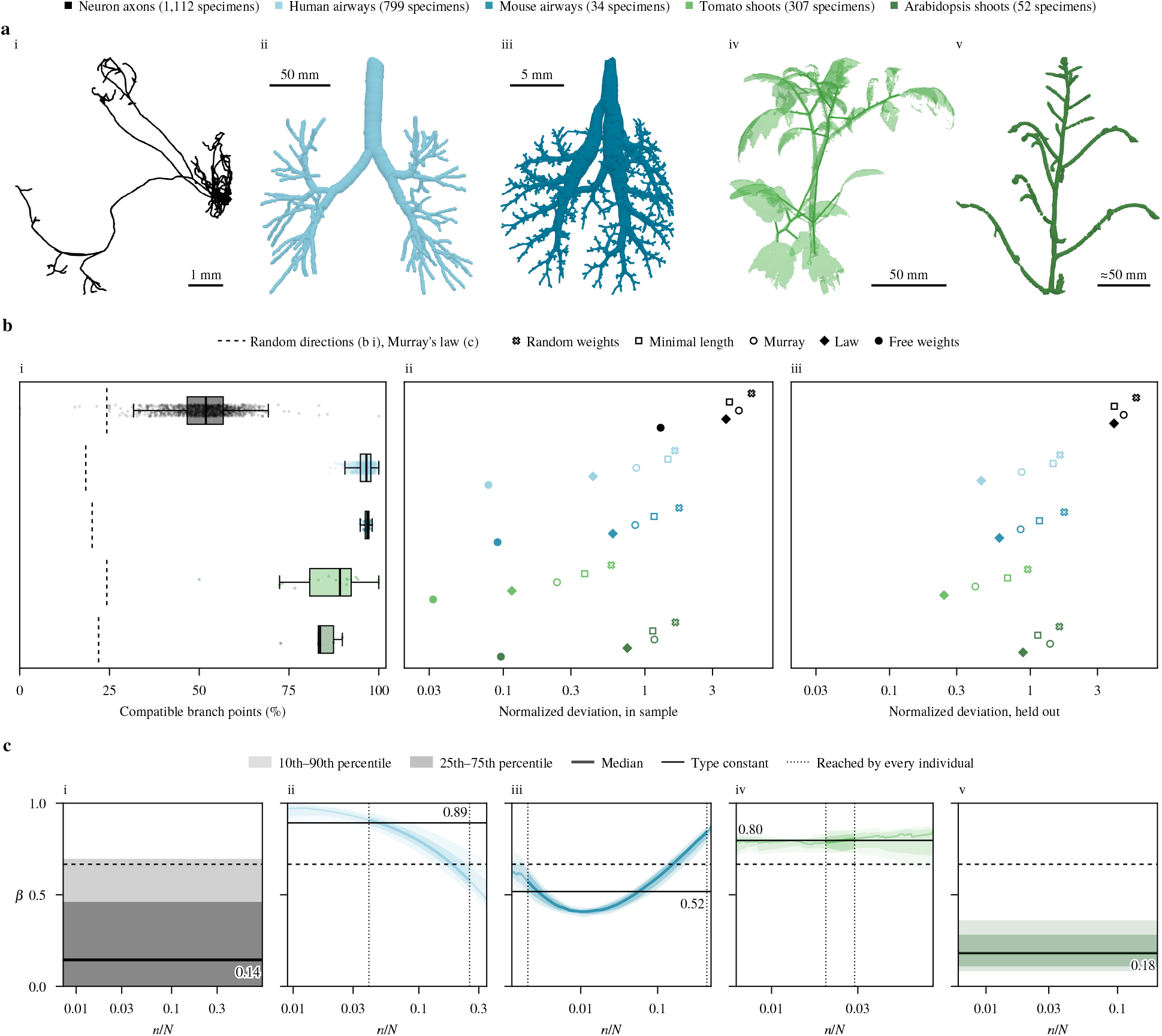
Airways, plant shoots and neuron axons. **a**, One specimen of each network (i–v); the Arabidopsis scale is approximate. **b**, (i) Share of compatible branch points per individual; random directions keep the observed vessel lengths. Boxes show quartiles and medians, and whiskers reach at most 1.5 interquartile ranges beyond the box. (ii) Median normalized deviation in sample under commonly assumed weightings (at *δ* = 0), the fitted law and free weights. (iii) As (ii), with each individual held out in turn. **c**, The fitted laws over the individuals against the terminal share *n/N* of a segment. Faded denotes where not every individual reaches. Deviations are normalized by each network’s median vessel length. *n* = 1,111 neurons (98 in b (ii), 97 for its law and in b (iii)), 799 patients, 34 mice, 17 tomato (219 specimens in b (ii), b (iii) and **c**) and 5 Arabidopsis plants (4 scanned on the same day in **c**).

The ladder rejects the weighted length for neuron axons, which fail its first two tests. Only about half of their branch points are compatible (Fig. 6b(i)), and their normalized deviation is up to 40 times higher than in the other four networks (Fig. 6b(ii)). The best constant law is minimal length, whose trees branch at a median 118^°^ where axons branch at 68^°^. Yet their type law beats Murray’s law in 73 of 97 axons (Fig. 6b(iii)), so the last two tests alone would have missed the failure.

Airways carry air instead of blood, so they test the weighted-length hypothesis with another fluid. In human lungs, the type law beats Murray’s law in 796 of 799 patients (Fig. 6b(iii)). The best constant *β* = 0.90 lies near Uylings’s limit for turbulent flow, *β* = 6*/*7 [12], yet beats it in 511 of 799 patients. Along the airways, *β* rises toward the terminals in 792 of 799 patients (Fig. 6c), consistent with measurements in bronchial casts [38]. Mouse lungs are as compatible as the human lungs (Fig. 6b(i)) but take *β* = 0.52 (Fig. 6c), and their *β* instead falls toward the terminals in all 34 mice. Cut to the depth of the human trees, this difference remains. Both curves are narrow, with an interquartile range of *β* over individuals of only 0.07 in humans and 0.05 in mice (Fig. 6c).

In plant shoots, a weight may stand for support, biomass or transport. Tomato shoots take *β* = 0.80 and branch at narrower angles than Murray’s law predicts. Conn et al. [11] proposed that these plants trade total length against the summed path length from the base to each terminal, a weighted length with weights 1 + *cn*_*v*_. With *c* fitted, it loses to the best constant in 15 of 17 plants. In Arabidopsis shoots, the minimal surface [17] (*β* = 1*/*3) beats Murray’s law in all five plants but the best constant (*β* = 0.24) in only two. The ladder thus rejects the weighted length where it fails, and elsewhere resolves laws fine enough to tell human from mouse lungs apart.

## Discussion

Optimality principles of branching networks, from vessels and airways to neurons and plants, have mostly been tested at single branchings, with new studies replacing one scaling parameter with another. The radius exponent of vascular trees, for example, changes with the estimator on one data set [19] and between coronary studies [16]. Meng et al. found that length minimization fails at branchings and predicts networks 25% shorter than measured [17]. By contrast, here we find that measured centerlines lie near the optimum of a weighted length, and that their geometry alone selects its scaling law. Our framework turns a design principle into a prediction of the whole network geometry, so that every branch point is compared directly with its measured position. Strict convexity gives each law exactly one global optimum, and dynamic programming finds the discrete optimum over the heart surface. The search recovers prescribed laws on synthetic trees, and every fitted law is judged on held-out trees.

Real vascular geometry places three of four vessel types above Murray’s law, and the law of the hepatic veins predicts the lung trees across organs and data sets. A larger *β* trades total length for shorter paths from the root, and narrows a symmetric branching from 75^°^ under Murray’s law to 55^°^ at *β* = 0.83. The pulmonary artery law (*β* = 0.83) corresponds to a radius exponent 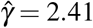, within the range of 2.1 to 2.6 measured in four species [40]. The portal vein stays near Murray’s law, and its law transfers to no other type. The radii of one human liver cast also suggest a lower radius exponent for the hepatic vein than the portal vein [41], which matches their fitted *β*, though no mechanism is known. The skeleton tells these two trees apart in nearly every liver, where the measured radii do not. Coronary trees on the heart surface lie near area preservation, which pulse waves may favor in large arteries [3].

Human and mouse airways are equally compatible yet take different laws. In mouse lungs, side branches bud in rows along a parent airway [42], which could explain their wider branchings (83^°^ against 58^°^ in humans) and lower *β* . Neuron axons fail the validation ladder, so the framework rejects a forward model where it does not hold.

A physiological reading of *β* needs radii, pressure and flow on the same trees. Models of structural adaptation [34] and synthetic trees optimized under a known cost [5, 6] could link *β* to a mechanism, and healthy cohorts could give the law for healthy vessels. Public dataset releases now hold thousands of networks, five of the six datasets used were released since 2025 (Supplementary Table 2). A design principle can therefore be tested on whole geometries, on held-out patients and as a law that varies along the tree. The framework takes any forward model solved to a global optimum (Eq. (2)) and any family of scaling laws, not only the weighted length.

Our study has two limitations. First, free weights describe measured geometry only in sample, and place held-out nodes worse than the law in nearly every tree (Supplementary Fig. 4). Type laws still have a clear gap to the free weights and the measured geometry. On trees of known law, simulated missing side branches and other artifacts reproduce most of this gap, which suggests that it comes from the scans rather than the model (Supplementary Fig. 6). The inferred *β* is robust to these artifacts, which shift it by at most 0.08 on average, under half the difference between hepatic and portal veins. Second, the geometry identifies which law places the vessels, not why, and 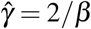 is a comparison under a volume cost, not a measured exponent.

## Online Methods

### Mathematical framework Trees

We describe each tree’s centerline as a directed graph *T* = (*V, A*) with nodes *u* ∈ *V* and segments *a* = *uv* ∈ *A*. Each segment runs from its proximal node *u* to its distal node *v*. The (single) *root* node 0 is the proximal node of the root segments, and each node without children is a *terminal node v* ∈ *L*?⊂ *V*. In every centerline, we remove all nodes with one parent and one child. Thus, a tree consists only of a root node, terminal nodes and branch nodes. The geometry of each tree is the position *x*_*u*_ of each node *u* and the length *l*_*a*_ = ∥*x*_*u*_− *x*_*v*_∥ of each segment.

We define the *root region V*_0_ ⊂ *V* as the root and the nodes of each vessel leaving it, up to where that vessel exits the root vessel’s radius hull. Together with the terminal nodes and the other nodes fixed during conversion, it forms the set of fixed nodes *F*, which keep their measured positions (*Datasets and trees*). The forward model places only the remaining free branch nodes *V* \ *F*.

The *perfusion territory n*_*v*_ = |*L*_*v*_| of node *v* is the number of terminal nodes *L*_*v*_ distal to it, so *n*_*v*_ = 1 ata terminal node and *N* = *n*_0_ counts all terminal nodes. We assume all terminals draw equal flow, so *n*_*v*_*/N* is the *flow fraction* of segment *a* = *uv*, the share of the root flow that enters the subtree distal to *v*.

### Inverse problem

A design principle is a family of *forward models F*(**x**; *θ*) with parameters *θ* ∈ Θ. The forward problem places the nodes **x** = (*x*_*v*_)_*v*∈*V*_ under one *θ* . An *outer search* selects the *θ* whose placement achieves the best *score D*. Thus, the inverse problem is

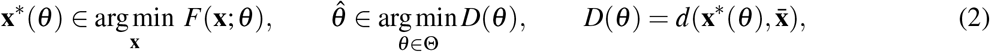

where the fixed nodes stay at the positions prescribed by their centerline 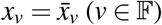 and *d* is the deviation of two geometries of the same tree topology. The solver of the forward problem must find a global optimum, so that the computed score *D* depends solely on *θ* . This holds when *F* is strictly convex, as the weighted-length model is for *δ >* 0. Each law defines one second-order cone program, solved to a certified global optimum in 14 ms on the largest tree, so no law is judged by a local optimum. The framework does not assume that trees attain the optimum but tests the geometry each law predicts.

Here, *F* is a weighted-length problem with *θ* defining the scaling law that maps the flow fraction of each segment to a weight. The score *D* measures the displacement averaged over the vessel length, and the outer search is a staged grid search followed by a pattern search.

### Weighted length model

Given a positive weight *w*_*a*_ for each segment, we cast the forward model into a nonlinear optimization problem (NLP). We include all variables to be optimized in the vector *y* = (**x**, *l*), with the nodal positions **x** = (*x*_*v*_)_*v*∈*V*_ and the segment lengths *l* = (*l*_*a*_)_*a*∈*A*_. The best geometry is then found in the set

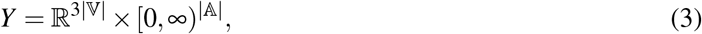

and our NLP reads:

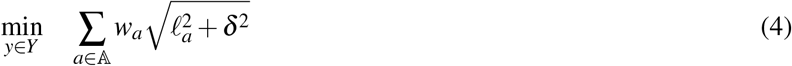

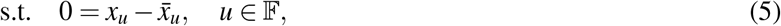

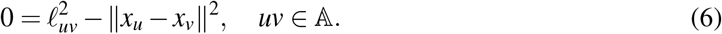

Here Eq. (5) keeps the fixed nodes at their measured centerline positions 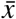, and Eq. (6) ensures consistency between nodal positions and segment lengths. *δ* ≥ 0 is the smoothing length, which keeps short vessels from shrinking to a point, and is set to *δ* = 5 mm for every vascular tree. At *δ* = 5 mm, the mean deviation of each vessel type lies within 1.5% of its lowest value over *δ* from 0 to 16 mm (Supplementary Fig. 1d). Each tree keeps its law of three control points fitted at 5 mm in this comparison. Between *δ* = 0, 5 and 8 mm, the best constant *β* of each type moves by at most 0.03 (Supplementary Fig. 1e).

For numerical efficiency, we write Eqs. (4) to (6) as a second-order cone program. Its variables *y* = (**x**,*t*) ∈ ℝ^3|*V*|^ ×ℝ^|*A*|^ replace each segment length by a bound *t*_*a*_ on the smoothed length, and the problem reads:

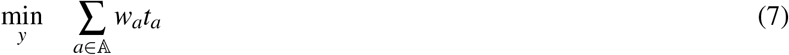

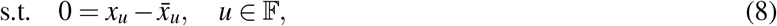

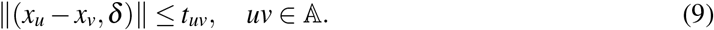

Eq. (8) fixes the same nodes as Eq. (5), and Eq. (9) ensures that *t*_*uv*_ bounds the smoothed length of segment *uv*, which it equals at the optimum since every weight is positive.

### Scaling laws

Each family of scaling laws has its own parameters *θ* . The family of scaling laws used here is defined by

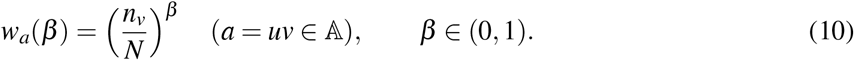

A constant exponent *β* covers the laws between the limits *β* = 0 (minimal length) and *β* = 1 (area preservation), with *β* = 2*/*3 giving Murray’s law. For one constant *β* per tree, *θ* = *β* .

To allow *β* to vary across the tree’s hierarchy, we describe its shape with a Bernstein curve of *M* control points

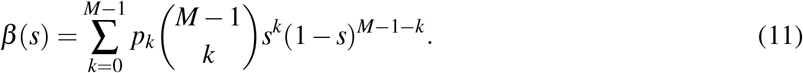

The control points are *θ* = (*p*_0_, …, *p*_*M*−1_) ∈ (0, 1)^*M*^. All network types take their respective *M* of Supplementary Table 1. Here *s* describes the position on the tree through the coordinate of segment *a* = *uv* with

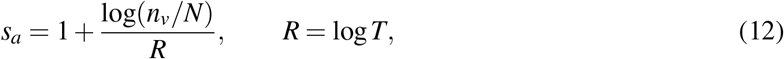

with *s* = 1 denoting the root and *s* = 0 denoting the terminal nodes of the pruned tree at flow fraction 1*/T* . Below *s* = 0, *β* stays at *p*_0_. The pruning of centerlines ensures that trees are fitted and compared fairly across the same depth, and the choice of *T* is given in *Datasets and trees*.

Lastly, weights are defined by integrating the exponent,

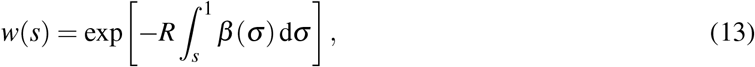

where *σ* runs over the tree coordinate from *s* to the root, and segment *a* takes the weight *w*_*a*_ = *w*(*s*_*a*_). This formulation gives *w* = 1 at the root and the constant law when all control points are equal. The scaling law ensures 0 *< β <* 1 by design and consequently all produced weights are positive and subadditive at every branch point.

If flow is proportional to territory, radius scales as 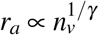 and the cost of a vessel is its volume 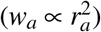.

We can then relate the exponent *β* to the radius exponent with

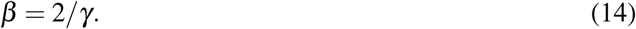

We report 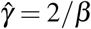 only under these assumptions, and to allow comparisons with measured radius exponents of the literature.

### Deviation

Predicted and measured trees share their topology, so their segments pair by identity. We use the Fréchet distance to measure the deviation between the predicted and measured chords of a segment. The deviation is its mean weighted by the measured segment length,

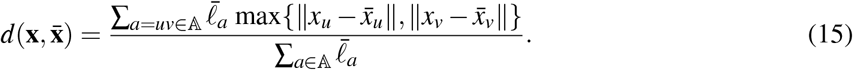

To compare trees of different size, we divide *d* by the median measured vessel length of their type or network, giving the *normalized deviation*. As a check, we also tried a different score function in place of *D*, the relative gap 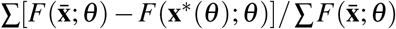, with sums over the trees of a type. The constant *β* it picks differs from the one *D* picks by 0.01 to 0.05 in the four vessel types, and by a median of 0.02 tree by tree (1,643 of 1,819 trees within 0.05).

### Outer search

For each tree, we seek the law with the smallest deviation score 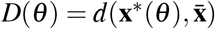. Our search is free of derivatives and has two phases.

The first phase is a staged grid search. For each control point, three evenly spaced values in a prescribed interval are tried. Each stage solves all combinations and keeps the one with the smallest score. The next stage halves the interval around the best found parameter values. The interval starts as [0, 1], over seven stages. For the constant law, the first stage tries nine values instead of three.

The second phase is a greedy pattern search from the best found grid point, with a step length *η* = 1*/*8. It steps each control point up and down by *η*, then each pair of control points along the four diagonals. It moves at once to any step that lowers *D* by more than 10^−6^ of its value. After a sweep without a move it halves *η*, and it stops once *η* falls below 1*/*256.

### Extension to surface-constrained problems

Coronary arteries run over the outside of the heart, so a straight segment can cut through the heart wall. We only need to replace the forward problem of the weighted-length model Eqs. (4) to (6) to use the framework on a surface. The tree, its fixed nodes, the weights *w*_*a*_, the smoothing length *δ* = 5 mm, the deviation Eq. (15) and the score *D* carry over from the *Mathematical framework*. Only two things change. First, each free branch point can now take only finitely many points on a surface, instead of any position in ℝ^3^. Second, the length *l*_*a*_ of a segment is the length of its shortest path over this surface, instead of its straight length in Eq. (6). The problem becomes discrete and the cone program does not apply. Instead, we solve it exactly by dynamic programming.

### Points and lengths on the surface

We start from a triangle mesh of the surface the vessels run on. We thin its vertices to a minimum spacing, and these points *P* are the places a free branch point may take. Every two points closer than a prescribed threshold are joined by a straight segment. The points and segments form a graph *G* = (*P, E*) over the surface. We write *l*_*G*_ (*p, q*) for the length of the shortest path between points *p, q* ∈ *P*. A segment between two free with such a fixed node first(runs straight to a near)by surface point *p* ∈ *P*, and then along the surface. Its length nodes has the length 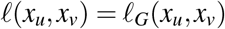. A fixed node *u* can lie off the surface at position 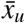. A segment is then 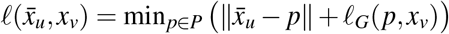.

### Surface forward problem

The forward problem then reads:

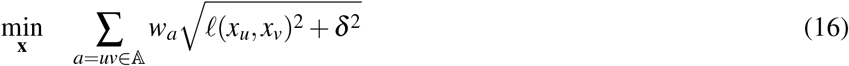

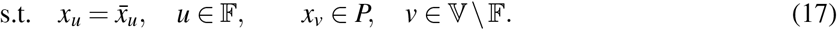

Eq. (16) is Eq. (4) with surface lengths, and Eq. (17) keeps the fixed nodes of Eq. (5) and puts every free node on a point of the surface.

### Exact solution by dynamic programming

Each free branch point chooses among finitely many points, so the problem is not convex. It is, however, a sum over the segments of a tree with fixed terminals, so we solve it exactly by the dynamic programming of Sankoff and Rousseau [22]. For a node *v* at point *x*, let *C*_*v*_(*x*) be the least cost of the subtree below *v*, with *C*_*v*_ = 0 at a terminal node. From the terminals to the root,

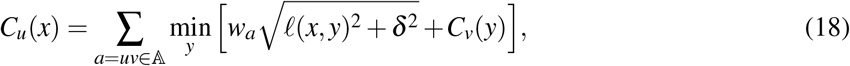

where *y* runs over *P* for a free child and is 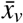 for a fixed one. 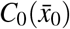 is the optimum of Eqs. (16) to (17), and the minimizing points are read back down the tree. As in the generalized distance transforms of Felzenszwalb and Huttenlocher [43], one Dijkstra search over the graph, extended to the smoothing length *δ*, gives the inner minimum of Eq. (18) for every point *x* of the parent at once. This step is exact over the points. Before solving, bounds on the cost discard the points no optimal placement can use, and on our test cases this kept every minimizing point.

### Datasets and trees

Vessel trees come from seven public releases of contrast-enhanced CT. All human data come from public, de-identified releases collected under the approvals of their original studies, so this study needed no new ethics approval. We use one tree per patient and vessel type. After pruning, 1,819 trees with 460,324 branch points remain.

Radii enter no fit and no forward model of a law, only the root region, the local branching analysis and the weights read from radii (Supplementary Fig. 7). The radius of a vessel is the median of the measured radii at the centerline points inside it, or at its two ends when it has none.

### Converting centerlines to trees

The radius hull of the root vessel is a tube of interpolated radius (a ball at a root with several children). We fix the root region because centerline extraction lays these nodes on the axis of the root vessel rather than where the vessels branch. The centerlines hold no loops, and branch points with more than two daughters stay as they are.

We prune each tree to the depth *T* its images resolve. We keep the first segment whose territory falls below *N/T* and remove the subtree below it. Its distal node becomes a fixed terminal with the territory of the removed subtree. A patient enters only if *N* ≥ *T* (Supplementary Table 2). The airways, plant shoots and axons are not pruned, and *R* is the logarithm of each network’s largest territory.

### Hepatic and portal veins

VEELA [23], LIRCAD [24] and HVA-CT [25] supply the hepatic and portal veins. We traced the liver centerlines with VMTK from roots picked by hand, which discards any part not connected to the root. A hepatic vein tree drains into the vena cava, which leaves the segmented liver at its lower border. We root the tree at the upper end of the vena cava, fix the path to its lower end, and count that end as an *exit terminal*. Its share of the whole territory is 0.6, the fraction of the flow at the upper end of the vena cava that comes from below the liver [44]. *N* is then the count of plain terminals divided by 0.4, and territories are real numbers. The prune depth is 100 for hepatic veins and 40 for portal veins. Portal veins fix the root region, the root and the terminals; hepatic veins also fix the path to the exit terminal.

### Pulmonary vessels

PTL [26] and HiPaS [27] supply the pulmonary arteries and veins, and PARSE [28] only arteries. PTL comes with centerlines, HiPaS and PARSE with the centerline graphs of VesselTok [45]. In the lungs, we also fix the path from the root to the first branch point whose two daughters each feed at least 0.1 of the tree’s terminals. The pulmonary vein graphs of PTL and HiPaS come with the vessels at the atrium marked as fixed, and we keep these marks. The prune depth is 200.

### Coronary arteries

We use the left coronary trees of ImageCAS-X [29, 30], with the root at the start of the centerline and the branch and end points as nodes. Only the root and the terminals are fixed, so *V*_0_ = {0} . For cases 1 to 50, the heart wall is the surface of the whole heart, with the atria, the aortic root and the great vessels. For cases 51 to 200, it is the surface of the left ventricular myocardium and both ventricles. All surfaces were segmented by TotalSegmentator [46], smoothed and remeshed with VMTK [47] to triangles with edges of about 1 mm.

For coronary trees, the surface forward problem is defined on the heart wall, with each vertex moved outward along its normal by 2.5 mm, the median signed distance of all trees’ roots and terminals from the wall. The minimum spacing is 1 mm and the threshold for joining points is 3.5 mm.

### Airways

We take the human airways of PTL and the mouse airways of lapdMouse, one tree per individual. We fix the trachea up to the first branch point whose two children each hold at least 10% of the root territory, and both main bronchi up to their next branch point. To cut the mouse lungs to the depth of the human trees, we keep their branch points up to generation 7, the human median. We also fix the root region, the root and the terminals, with no other preprocessing.

### Plant shoots

We use the tomato shoots of Conn et al. [11] and the Arabidopsis shoots of Meng et al. [17] (Supplementary Table 2). We fix the root region, the root and the terminals, with no other preprocessing.

### Neurons

We use the MouseLight axons (Supplementary Table 2). We fix the root region, the root and the terminals, with no other preprocessing. Every other branch point is free.

### Analyses

#### Local branching analysis

We read every centerline file of the four vessel types before pruning and 86 coronary trees traced from the original ImageCAS labels, 2,559 trees in all. We measure every branch point with two daughters off the fixed nodes, 563,088 in total. At a branch point with parent radius *r*_1_ and daughter radii *r*_2_ and *r*_3_, we solve Eq. (1) for *γ* when both daughters are thinner than the parent, and otherwise no positive *γ* exists. Murray’s law predicts the angle 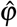 between the daughters by

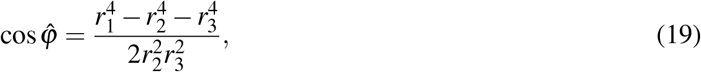

and we measure the angle *ϕ* between the two daughter chords. Strahler orders are counted on the full centerline tree.

#### Compatible branchings

At the optimum of Eqs. (4) to (6) with *δ* = 0, the unit vectors from a free branch point to its neighbors, multiplied by their positive weights, sum to zero. Such weights exist only when the branch point lies inside the convex hull of its neighbors. At a branch point with two daughters, we project the branch point onto the plane of its three neighbors. It is compatible when its projection lies inside their triangle and its height above the plane is below 0.2 of the mean length of its three segments. At a branch point with three or more daughters, we test whether the origin lies in the convex hull of the directions to its neighbors. We test noise by moving every free node by Gaussian noise of root mean square 0.5, 1, 2 and 4 mm.

#### Free weights

Free weights give each segment with a free end its own weight. Starting from the tree’s fitted law of three control points, we fit the logarithms of the weights to the free-node positions by least squares with the Gauss–Newton method, keeping *δ* = 5 mm. To test them on held-out nodes, we fit them on every other free node and read the root-mean-square displacement of the remaining free nodes.

#### Recovery of prescribed laws

To validate the inference, we solve measured topologies under a prescribed law, add noise, and fit the result as if it were measured. From each vessel type, we take 75 trees evenly spaced in size and assign them in turn to three prescribed laws. The three laws are Murray’s law, a law whose control points rise evenly from 0.35 to 0.80, and a cup shape with control points 0.85, 0.25 and 0.85. We solve each tree under its prescribed law at *δ* = 5 mm and add Gaussian noise of standard deviation 0.25 mm to every coordinate of every node, fixed nodes included, and fit its law of three control points. For coronary trees, we prescribe constant *β* of 0.67, 0.83, 0.86, 0.90 and 0.97 on 50 trees evenly spaced in size, and fit each alone under a constant law.

### Held-out test and transfer

The best constant *β* of a type is the one that minimizes the mean deviation over its trees (Supplementary Fig. 8a). The type constant and type law of a held-out tree are the medians of the constant *β* and of *β* (*s*) over all other trees of its type. The median is computed at 281 equally spaced points over *s* in [0, 1]. Each vessel type takes its number of control points from Supplementary Table 1, and portal veins take three in Fig. 4. For transfer, we apply the law of a whole type to every tree of another, with the *R* of the receiving type.

For coronary trees, we fit each of the 151 trees alone and test each under the median *β* of the other 150. Normalized deviations use 25.87 mm, the median vessel length of all 151 trees.

For the airways, plant shoots and axons, each network takes its own *δ*, defined at the best constant *β* . Its type constant and type law are defined like the vessel types. Each network takes the number of control points that its held-out individuals support (Supplementary Table 1). Starting from one, we add a control point while it lowers the held-out deviation by at least 0.1% on average, with a 95% bootstrap interval above zero, and stop at four. We chose this floor after seeing the gains, and any floor from 0.04% to 0.18% gives the same numbers. The law of Conn et al. weights a segment by 1 + *cn*_*v*_, with *c* chosen on the other plants.

#### Statistics

We analyze a branch point during local analysis and a global tree geometry otherwise. Two laws are compared by their score *D* on the same trees. Sign tests are exact and two-sided, and we report *p* values without correction for multiple comparisons. The interval of the best constant *β* of a type is the 2.5th to 97.5th percentile over 10,000 bootstrap draws of patients within the type, each with its own best constant. Constants within 1% of a type’s lowest mean deviation span at most 0.04 in each of the three types and 0.08 in portal veins (Supplementary Fig. 1e). The effects on held-out trees use 10,000 draws over patients, and the comparison of the datasets of one type draws within each dataset (Supplementary Fig. 5). No statistical method was used to predetermine sample size.

### Computation

Our inference framework runs on Julia 1.13.1 [48]. We solve the cone program Eq. (7) with the interior-point solver Clarabel [49] to a certified optimum at a tolerance of 10^−9^. When improved computational speed is needed (free weights, held-out nodes), we use Newton’s method with Armijo backtracking instead. It stops once no variable moves by more than 10^−8^ mm. We run 3D Slicer 5.12.4 [50] with the extensions VMTK [47] and TotalSegmentator 2.14.0 [46]. Fitting all 1,819 vascular trees took 2.7 hours. All timings reported here come from an Apple MacBook Pro with an M4 Max chip and 36 GB of memory.

### Use of artificial intelligence

OpenAI’s GPT-5.6 Sol and Anthropic’s Claude Opus 5.5 were used during manuscript preparation for writing and language refinement. Claude Opus 5.5 was used for limited code assistance and debugging. All scientific content, computational methods, code and results were reviewed and verified by the authors.

## Data availability

All trees come from public datasets: the hepatic and portal veins from VEELA [23], LIRCAD [24] and HVA-CT [25]; the pulmonary arteries, veins and human airways from PTL [26], HiPaS [27] and PARSE [28], the latter two as the centerline graphs of VesselTok [45]; the coronary arteries from ImageCAS [29] and ImageCAS-X [30]; the mouse airways from lapdMouse [31]; the plant shoots from Conn et al. [11] and Meng et al. [17]; and the MouseLight axons [32] from NeuroMorpho.Org [51]. Supplementary Table 2 lists each dataset’s license. The liver centerlines we traced, the coronary centerlines traced from the ImageCAS labels and the tables behind every figure and number of this paper are provided to the editors and reviewers as a zip-file for peer review. They will be deposited on Zenodo on publication, each under the license of its source dataset.

## Code availability

BranchingLaws.jl, the Julia package that solves the forward models and fits the scaling laws, is provided to the editors and reviewers as a zip-file for peer review. It holds the converter of each dataset whose centerlines we did not trace, the scripts that write the tables behind every figure and number, and the scripts that plot the statistics panels. On publication, it will be made public under the MIT license at https://github.com/etiennejessen/BranchingLaws.jl and archived on Zenodo.

## Acknowledgment

The results presented in this paper were obtained as part of the ERC Starting Grant project “ImageToSim” that has received funding from the European Research Council (ERC) under the European Union’s Horizon 2020 research and innovation programme (Grant agreement No. 759001). The authors gratefully acknowledge this support.

## References

[1] Murray, C. D. The physiological principle of minimum work applied to the angle of branching of arteries. The Journal of General Physiology 9, 835–841 (1926).

[2] Murray, C. D. The physiological principle of minimum work: I. The vascular system and the cost of blood volume. Proceedings of the National Academy of Sciences 12, 207–214 (1926).

[3] West, G. B., Brown, J. H. & Enquist, B. J. A general model for the origin of allometric scaling laws in biology. Science 276, 122–126 (1997).

[4] Banavar, J. R., Maritan, A. & Rinaldo, A. Size and form in efficient transportation networks. Nature 399, 130–132 (1999).

[5] Jessen, E., Steinbach, M. C., Debbaut, C. & Schillinger, D. Rigorous mathematical optimization of synthetic hepatic vascular trees. Journal of The Royal Society Interface 19, 20220087 (2022).

[6] Jessen, E., Steinbach, M. C., Debbaut, C. & Schillinger, D. Branching exponents of synthetic vascular trees under different optimality principles. IEEE Transactions on Biomedical Engineering 71, 1345–1354 (2024).

[7] Jessen, E., Steinbach, M. C. & Schillinger, D. Optimizing non-intersecting synthetic vascular trees in nonconvex organs. IEEE Transactions on Biomedical Engineering 72, 2871–2881 (2025).

[8] Rinaldo, A., Rigon, R., Banavar, J. R., Maritan, A. & Rodriguez-Iturbe, I. Evolution and selection of river networks: Statics, dynamics, and complexity. Proceedings of the National Academy of Sciences 111, 2417–2424 (2014).

[9] Cuntz, H., Forstner, F., Borst, A. & Häusser, M. One rule to grow them all: A general theory of neuronal branching and its practical application. PLOS Computational Biology 6, e1000877 (2010).

[10] Chandrasekhar, A. & Navlakha, S. Neural arbors are Pareto optimal. Proceedings of the Royal Society B: Biological Sciences 286, 20182727 (2019).

[11] Conn, A., Pedmale, U. V., Chory, J. & Navlakha, S. High-resolution laser scanning reveals plant architectures that reflect universal network design principles. Cell Systems 5, 53–62.e3 (2017).

[12] Uylings, H. B. M. Optimization of diameters and bifurcation angles in lung and vascular tree structures. Bulletin of Mathematical Biology 39, 509–520 (1977).

[13] Savage, V. M., Deeds, E. J. & Fontana, W. Sizing up allometric scaling theory. PLOS Computational Biology 4, e1000171 (2008).

[14] Eloy, C. Leonardo’s rule, self-similarity, and wind-induced stresses in trees. Physical Review Letters 107, 258101 (2011).

[15] Huo, Y. & Kassab, G. S. Intraspecific scaling laws of vascular trees. Journal of The Royal Society Interface 9, 190–200 (2011).

[16] Taylor, D. J. et al. Systematic review and meta-analysis of Murray’s law in the coronary arterial circulation. American Journal of Physiology-Heart and Circulatory Physiology 327, H182–H190 (2024).

[17] Meng, X., Piazza, B., Both, C., Barzel, B. & Barabási, A.-L. Surface optimization governs the local design of physical networks. Nature 649, 315–322 (2026).

[18] Kassab, G. S. Scaling laws of vascular trees: Of form and function. American Journal of Physiology-Heart and Circulatory Physiology 290, H894–H903 (2006).

[19] Newberry, M. G., Ennis, D. B. & Savage, V. M. Testing foundations of biological scaling theory using automated measurements of vascular networks. PLOS Computational Biology 11, e1004455 (2015).

[20] Lippmann, P., Sanmartín, E. F. & Hamprecht, F. A. Theory and approximate solvers for branched optimal transport with multiple sources. In Advances in Neural Information Processing Systems, vol. 35, 267–279 (Curran Associates, 2022).

[21] Aswani, A., Shen, Z.-J. M. & Siddiq, A. Inverse optimization with noisy data. Operations Research 66, 870–892 (2018).

[22] Sankoff, D. & Rousseau, P. Locating the vertices of a Steiner tree in an arbitrary metric space. Mathematical Programming 9, 240–246 (1975).

[23] Yazıcı, Z. A. et al. VEELA: A clinically-constrained benchmark for liver vessel segmentation in computed tomography angiography (2026). Preprint at https://arxiv.org/abs/2605.22357.

[24] Machta, Y. et al. Enhancing the automatic segmentation and analysis of 3D liver vasculature models (2025). Preprint at https://arxiv.org/abs/2411.15778.

[25] Salvi, M. et al. HVA-CT: Hepatic venous annotations CT. Zenodo, 10.5281/zenodo.19850108 (2026).

[26] Xie, K., Yang, J., Wei, D., Weng, Z. & Fua, P. Efficient anatomical labeling of pulmonary tree structures via deep point-graph representation-based implicit fields. Medical Image Analysis 99, 103367 (2025).

[27] Chu, Y. et al. Deep learning-driven pulmonary artery and vein segmentation reveals demography-associated vasculature anatomical differences. Nature Communications 16, 2262 (2025).

[28] Luo, G. et al. Efficient automatic segmentation for multi-level pulmonary arteries: The PARSE challenge (2023). Preprint at https://arxiv.org/abs/2304.03708.

[29] Zeng, A. et al. ImageCAS: A large-scale dataset and benchmark for coronary artery segmentation based on computed tomography angiography images. Computerized Medical Imaging and Graphics 109, 102287 (2023).

[30] Bransby, K. M. et al. ImageCAS-X: A dataset and benchmark for coronary artery segmentation and centerline extraction in coronary CT angiography (2026). Preprint at https://arxiv.org/abs/2608.30404.

[31] Bauer, C., Krueger, M., Lamm, W. J. E., Glenny, R. W. & Beichel, R. R. lapdMouse: Associating lung anatomy with local particle deposition in mice. Journal of Applied Physiology 128, 309–323 (2020).

[32] Winnubst, J. et al. Reconstruction of 1,000 projection neurons reveals new cell types and organization of long-range connectivity in the mouse brain. Cell 179, 268–281.e13 (2019).

[33] Budd, J. M. L. et al. Neocortical axon arbors trade-off material and conduction delay conservation. PLOS Computational Biology 6, e1000711 (2010).

[34] Pries, A. R., Secomb, T. W. & Gaehtgens, P. Structural adaptation and stability of microvascular networks: Theory and simulations. American Journal of Physiology-Heart and Circulatory Physiology 275, H349–H360 (1998).

[35] Kamiya, A. & Togawa, T. Adaptive regulation of wall shear stress to flow change in the canine carotid artery. American Journal of Physiology-Heart and Circulatory Physiology 239, H14–H21 (1980).

[36] Zarins, C. K., Zatina, M. A., Giddens, D. P., Ku, D. N. & Glagov, S. Shear stress regulation of artery lumen diameter in experimental atherogenesis. Journal of Vascular Surgery 5, 413–420 (1987).

[37] Ronellenfitsch, H. & Katifori, E. Global optimization, local adaptation, and the role of growth in distribution networks. Physical Review Letters 117, 138301 (2016).

[38] Restrepo, J. G., Ott, E. & Hunt, B. R. Scale dependence of branching in arterial and bronchial trees. Physical Review Letters 96, 128101 (2006).

[39] Changizi, M. A. & Cherniak, C. Modeling the large-scale geometry of human coronary arteries. Canadian Journal of Physiology and Pharmacology 78, 603–611 (2000).

[40] Altieri Correa, S., Kachabi, A., Colebank, M. J., Miles, C. E. & Chesler, N. C. Revisiting murray’s law in pulmonary arteries: Exploring branching patterns and principles. Journal of Biomechanical Engineering 147, 081006 (2025).

[41] Debbaut, C. et al. Analyzing the human liver vascular architecture by combining vascular corrosion casting and micro-CT scanning: A feasibility study. Journal of Anatomy 224, 509–517 (2014).

[42] Metzger, R. J., Klein, O. D., Martin, G. R. & Krasnow, M. A. The branching programme of mouse lung development. Nature 453, 745–750 (2008).

[43] Felzenszwalb, P. F. & Huttenlocher, D. P. Pictorial structures for object recognition. International Journal of Computer Vision 61, 55–79 (2005).

[44] Muthusami, P. et al. Splanchnic, thoracoabdominal, and cerebral blood flow volumes in healthy children and young adults in fasting and postprandial states: Determining reference ranges by using phase-contrast MR imaging. Radiology 285, 231–241 (2017).

[45] Prabhakar, C. et al. VesselTok: Tokenizing vessel-like 3D biomedical graph representations for reconstruction and generation. In Computer Vision – ECCV 2026, 131–149 (LNCS, 2026).

[46] Wasserthal, J. et al. TotalSegmentator: Robust segmentation of 104 anatomic structures in CT images. Radiology: Artificial Intelligence 5, e230024 (2023).

[47] Antiga, L. et al. An image-based modeling framework for patient-specific computational hemodynamics. Medical & Biological Engineering & Computing 46, 1097–1112 (2008).

[48] Bezanson, J., Edelman, A., Karpinski, S. & Shah, V. B. Julia: A fresh approach to numerical computing. SIAM Review 59, 65–98 (2017).

[49] Goulart, P. J. & Chen, Y. Clarabel: An interior-point solver for conic programs with quadratic objectives. Mathematical Programming Computation (2026). 10.1007/s12532-026-00320-7.

[50] Fedorov, A. et al. 3D Slicer as an image computing platform for the Quantitative Imaging Network. Magnetic Resonance Imaging 30, 1323–1341 (2012).

[51] Tecuatl, C., Ljungquist, B. & Ascoli, G. A. Accelerating the continuous community sharing of digital neuromorphology data. FASEB BioAdvances 6, 207–221 (2024).

