## Supplementary material for "Inferring scaling laws of branching networks from their geometry by global optimization"

### 1 Supplementary material

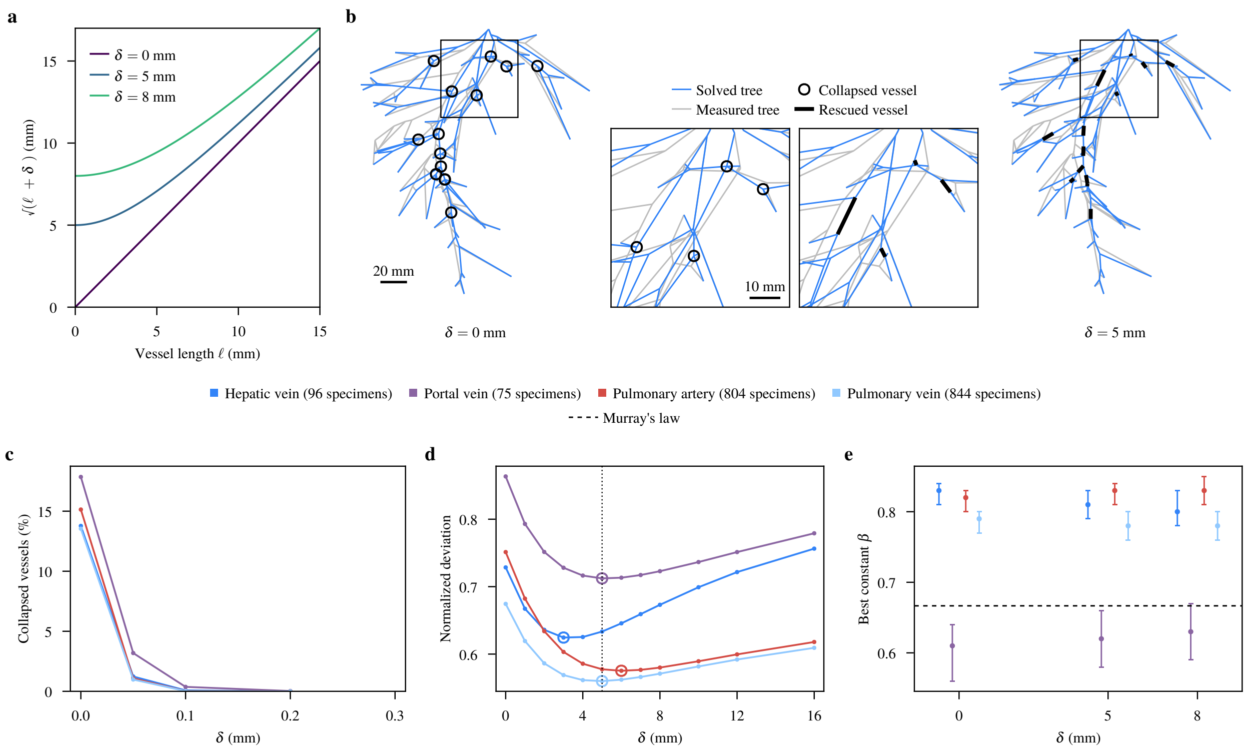


Supplementary Figure 1: **Collapsed vessels and the parameter $\delta$. a,** Cost $\sqrt{\ell^2+\delta^2}$ of a vessel of unit weight against its length $\ell$, for three values of $\delta$. **b,** A hepatic vein solved under its fitted law at $\delta = 0$ and $\delta = 5$ mm over its measured skeleton, with the boxed window magnified between them. A vessel collapses when its two branch points lie closer than 0.1 mm. **c,** Share of the vessels between two branch points that collapse, against $\delta$, pooled per type. **d,** Normalized deviation against $\delta$ under the laws fitted at 5 mm (dotted line); rings mark the lowest deviation of each type. **e,** Best constant $\beta$ of each type at three values of $\delta$, with the range of constants within 1% of its lowest mean deviation.

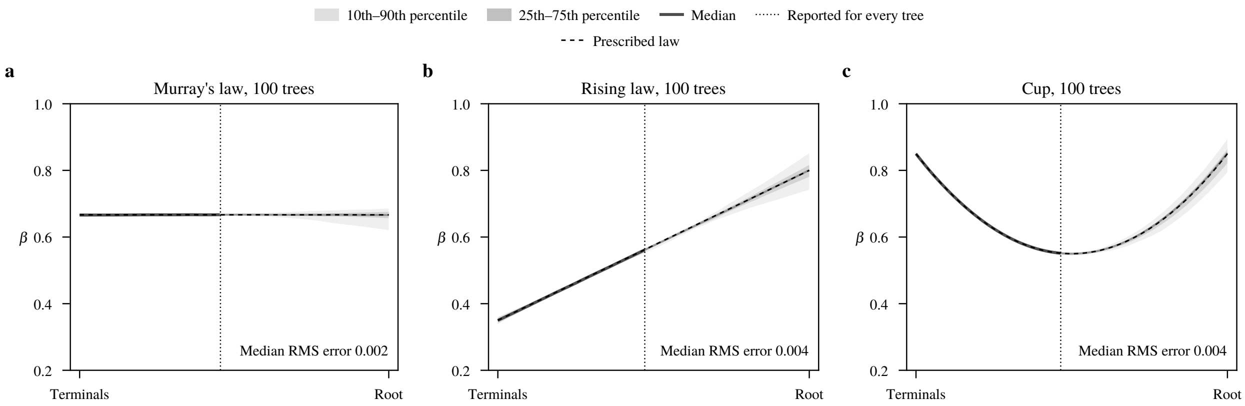


Supplementary Figure 2: **Recovery of prescribed laws. a–c,** Laws fitted to synthetic trees solved forward on measured topologies under Murray's law (**a**), a rising law (**b**) and a cup shape (**c**). Every node is moved by Gaussian noise of 0.25 mm standard deviation; 100 trees per law, 25 of each vessel type. Percentiles and median of the fitted exponent $\beta$ are taken over the trees. Left of the dotted line every tree has at least four free nodes at or above that point; right of it the bands are faded. The bands fan out towards the root because a tree has only one or a few free nodes there, so the noise on these few positions sets $\beta$. Without noise, the fit returns the prescribed law everywhere.

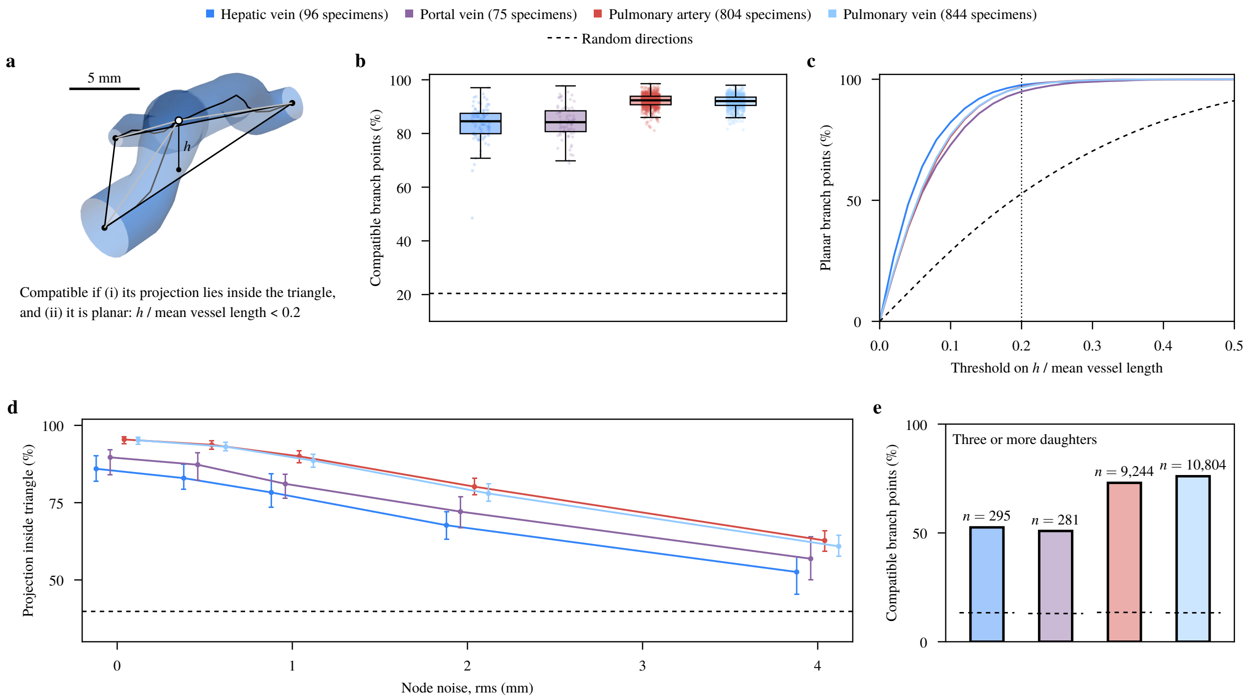


Supplementary Figure 3: **Compatibility of measured branch points. a,** A hepatic vein branch point (white) and the triangle of its three neighbours, over a render of its vessels; $h$ is its distance from the triangle's plane. **b,** Share of compatible branch points per specimen. Boxes show quartiles and medians, and whiskers reach at most 1.5 interquartile ranges beyond the box. **c,** Share of planar branch points against the threshold on $h$ over the mean vessel length; the dotted line marks the threshold of **a**. **d,** Share of branch points whose projection lies inside the triangle under random noise, median and quartiles over specimens. Node shifts of 2 mm root mean square lower the median share from 86.0–95.4% to 68–80%, and shifts of 4 mm to 53–63%. **e,** Share of branch points with three or more daughters at which the vessel directions hold the center in their convex hull, the condition of **a** for more vessels; $n$ gives the branch points. Random directions keep their observed vessel lengths.

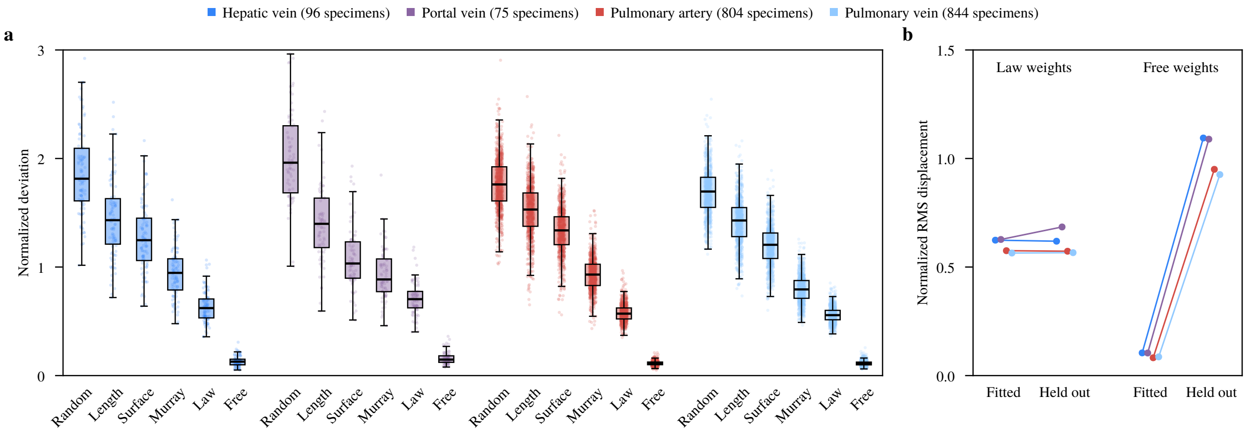


Supplementary Figure 4: **Commonly assumed weightings and free weights.** **a,** Normalized deviation per specimen under random weights (Murray weights shuffled among the vessels), minimal length (equal weights), minimal surface ($\beta = 1/3$) and Murray's law, all at $\delta = 0$, and under the fitted law and free weights, one positive weight per vessel, both fitted in sample at $\delta = 5$ mm. At $\delta = 5$ mm, Murray's law scores above the fitted law in all 1,819 trees, by a median of 9 to 38% per type. Boxes show quartiles and medians, and whiskers reach at most 1.5 interquartile ranges beyond the box. **b,** Displacement of the nodes the weights were fitted on and of the held-out nodes, every other free node, under the law and under free weights fitted to the other free nodes; points show the median of each type.

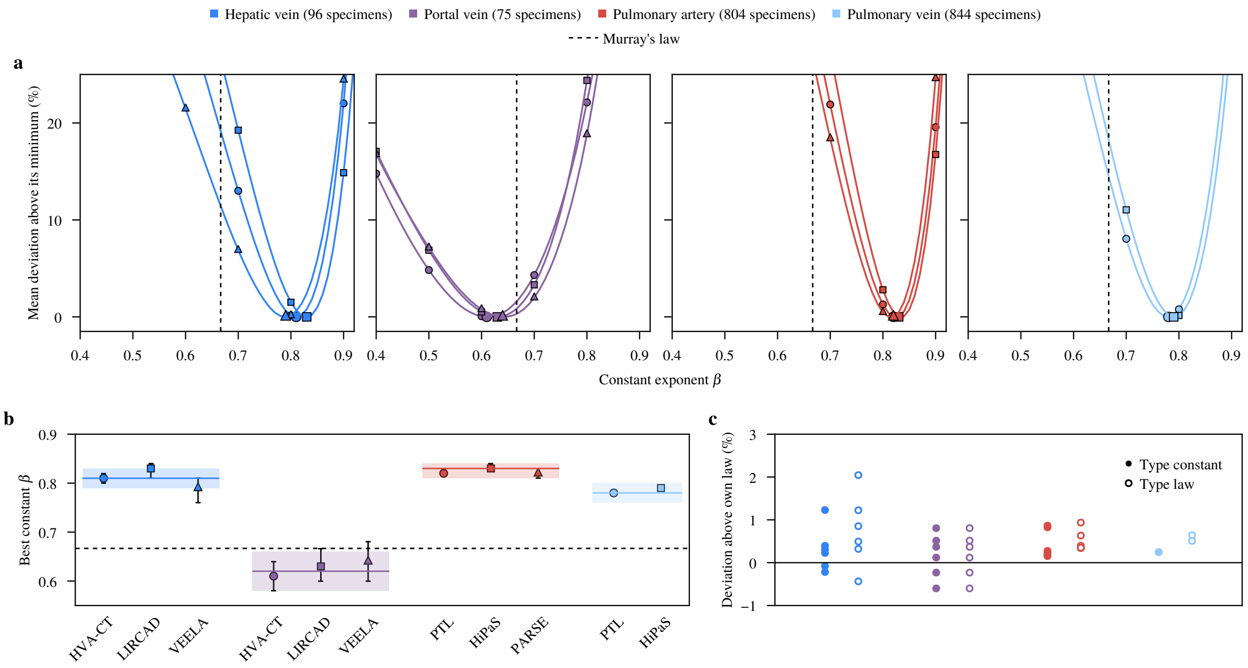


Supplementary Figure 5: **Agreement between the datasets of each vessel type. a,** Mean deviation against the constant exponent $\beta$ per dataset, in percent above the dataset's minimum, one panel per vessel type; markers tell the datasets apart. **b,** Best constant $\beta$ per dataset with its 95% interval over a bootstrap of patients; bands show the pooled best $\beta$ of each type and the range within 1% of its minimum. **c,** Mean deviation of the trees of one dataset under the law fitted on another dataset of their type, in percent above the law of their own dataset, for the type constant and type law; one point per pair of datasets.

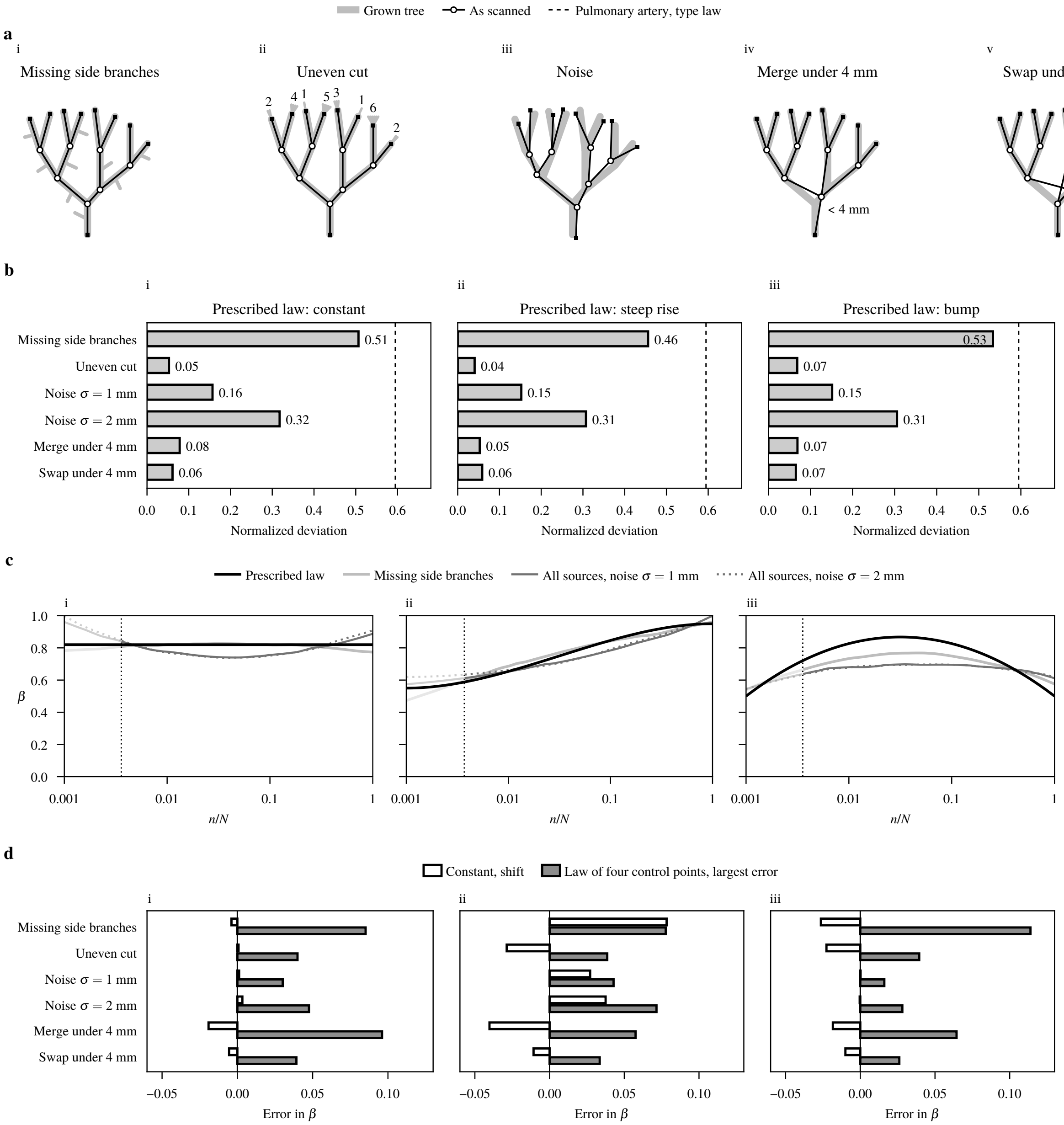


Supplementary Figure 6: **Deviation left under the prescribed law.** Trees grown under a prescribed law and cut to the vessels a scan resolves. **a,** Each source of error on a small tree. **i,** Side branches below the cut are missing. **ii,** The subtrees beyond an uneven cut hold different numbers of terminals, and each terminal is given their mean, 3. **iii,** Nodes carry noise. **iv,** Branch points closer than 4 mm merge. **v,** Branch points closer than 4 mm swap their order. **b,** Normalized deviation against prescribed laws under each above error source. **i,** Constant law. **ii,** Rising law. **iii,** Bump shaped law. Bars give the mean over ten grown trees. **c,** Fitted laws against prescribed laws. **d,** Shift of the fitted constant and largest error of the law (four control points) under each error source. Bars give the mean over ten grown trees.

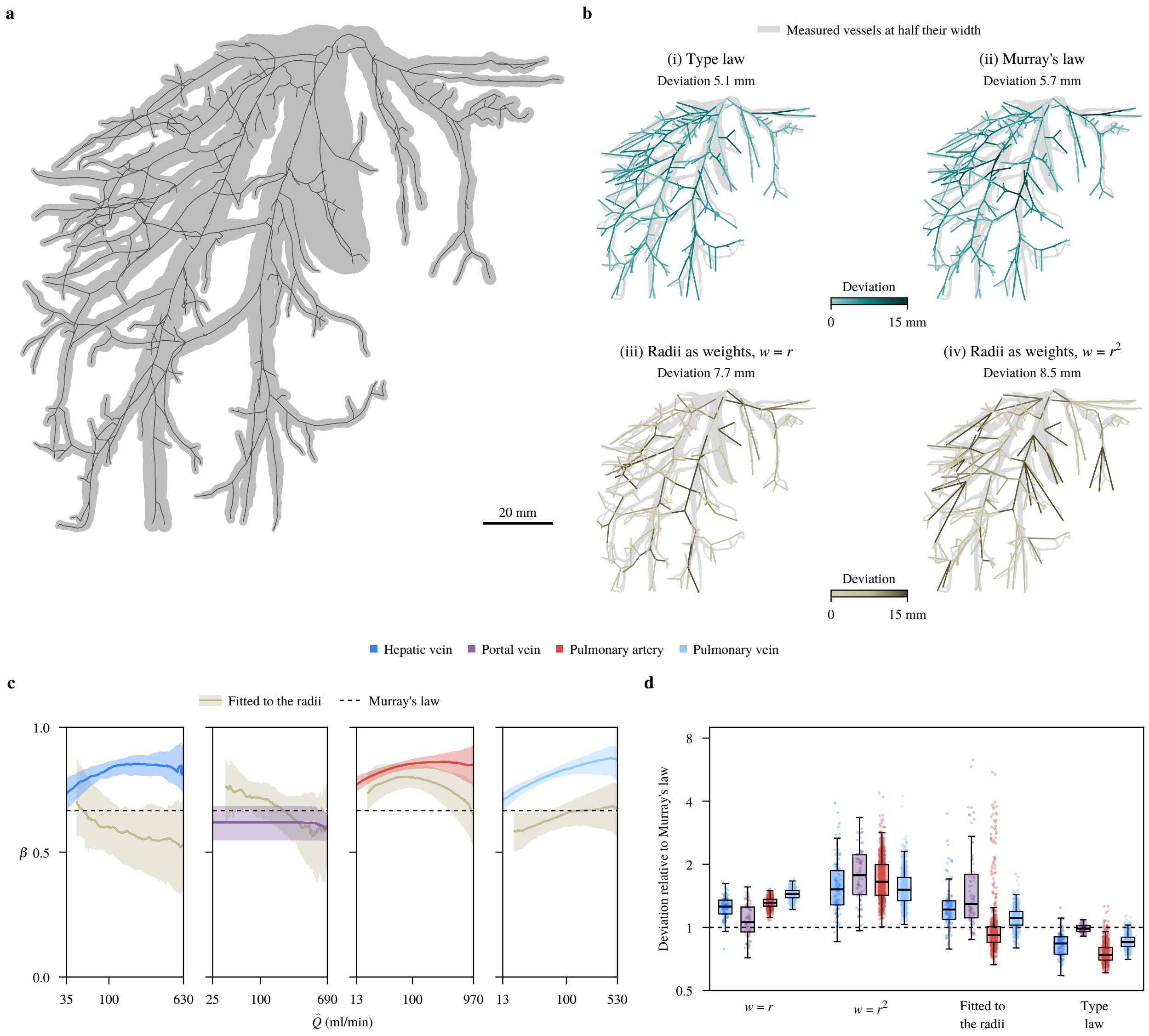


Supplementary Figure 7: **Weights from measured radii. a,** An unpruned hepatic vein (206 terminals), drawn at its measured diameter along its centerline. **b,** The tree of **a** solved under four weightings. Vessels are shaded by the larger deviation of their two ends, and titles give the deviation of the tree. (i) and (ii) use $\delta = 5$ mm, (iii) and (iv) $\delta = 0$, as a generation model would. **c,** Fitted laws of each type and laws fitted to the radii against the estimated flow $\hat{Q}$, median and 25th–75th percentiles where at least half of the trees reach, as in Fig. 3c. **d,** Deviation per tree relative to Murray's law at the same $\delta$ under $w = r$, $w = r^2$, the law fitted to the radii and the type law. Boxes show quartiles and medians, and whiskers reach at most 1.5 interquartile ranges beyond the box. The 109 pulmonary arteries and 60 pulmonary veins with a vessel of zero radius are excluded ($n = 96$, 75, 695 and 784 trees).

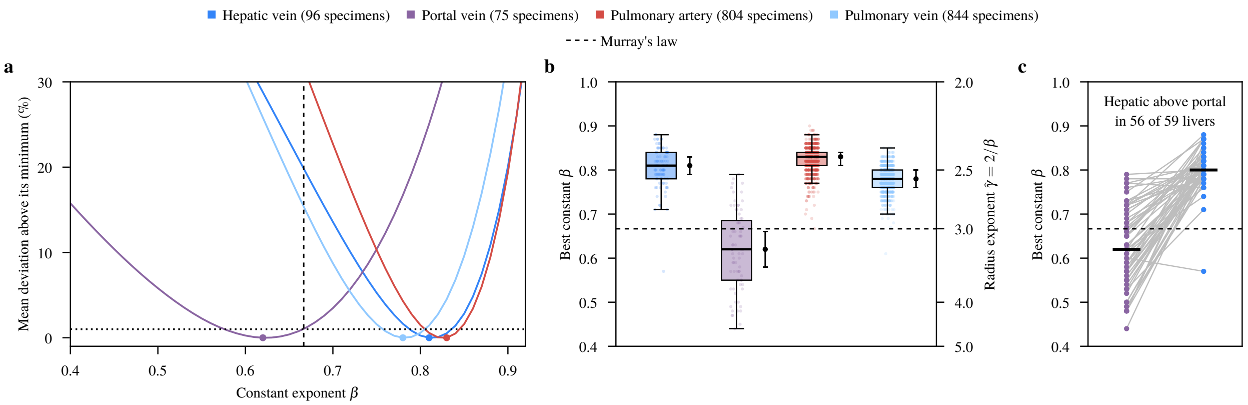


Supplementary Figure 8: **Best constant exponent of each vessel type. a,** Mean deviation over the specimens of each type against the constant exponent $\beta$, in percent above its minimum (dots); the dotted line marks 1%. **b,** Best constant $\beta$ per specimen, and the best $\beta$ of each type with the range within 1% of its minimum (black). Boxes show quartiles and medians, and whiskers reach at most 1.5 interquartile ranges beyond the box. **c,** Best constant $\beta$ of the portal and the hepatic vein of each liver that has both, joined by grey lines; bars mark the medians.

Supplementary Table 1: **Number of control points of each network.** Every tree is scored under its type law of one to four control points, the median of the fitted laws of the other individuals' trees (patients, mice, plants or neurons). The vessels are solved at $\delta = 5\,\text{mm}$, as in Fig. 4 of the main text, and the other networks at their own $\delta$, as in Fig. 6b (iii). Each column gives one step: the percent of the deviation that one more control point takes off, averaged over individuals, with its 95% bootstrap interval over individuals (10,000 draws). An individual's deviation is the mean over its specimens. A step counts (bold) when it takes off at least 0.1% on average and its interval lies above zero. Each network keeps the number before its first step that does not count, and the count stops at four. Every figure takes these numbers, except Fig. 4b(ii) and c, where portal veins take three. Any floor from 0.04% to 0.18% gives the same numbers, and we chose the floor after seeing the gains. The axons are the 100 drawn at random for Fig. 6, minus the three the solver failed on.

| Network | Individuals | One to two | Two to three | Three to four | Chosen |
|---|---|---|---|---|---|
| Hepatic vein | 96 | **0.78 [0.34, 1.22]** | **0.41 [0.26, 0.56]** | **0.35 [0.18, 0.52]** | 4 |
| Portal vein | 75 | 0.03 [−0.26, 0.32] | 0.10 [−0.11, 0.31] | 0.10 [−0.02, 0.21] | 1 |
| Pulmonary artery | 804 | **1.14 [1.01, 1.26]** | **0.24 [0.20, 0.27]** | 0.03 [0.01, 0.04] | 3 |
| Pulmonary vein | 844 | **1.78 [1.67, 1.89]** | **0.18 [0.15, 0.21]** | 0.04 [0.03, 0.05] | 3 |
| Human airway | 799 | **2.25 [2.10, 2.39]** | **1.29 [1.17, 1.42]** | **0.87 [0.82, 0.93]** | 4 |
| Mouse airway | 34 | **3.29 [2.53, 4.02]** | **5.08 [4.57, 5.57]** | **0.91 [0.50, 1.31]** | 4 |
| Tomato shoot | 17 | **2.98 [0.03, 6.13]** | −0.41 [−1.42, 0.61] | −1.78 [−2.97, −0.58] | 2 |
| Arabidopsis shoot | 5 | 0.44 [−0.09, 1.07] | 0.85 [−0.44, 1.89] | −0.60 [−1.86, 0.22] | 1 |
| Neuron axon | 97 | −1.30 [−2.29, −0.34] | −0.43 [−1.61, 0.53] | 0.33 [−0.32, 0.87] | 1 |

Supplementary Table 2: **Datasets.** Cases of each release that we used, and specimens the trees we fitted, one per patient. A case gives no specimen when its centerline could not be traced, or when its tree has fewer terminals than the prune depth: 100 for the hepatic vein, 40 for the portal vein and 200 in the lungs. Terminals per specimen are the median (range) after pruning, and branch points the total over the specimens. The hepatic and portal veins of one liver dataset come from the same patients, and so do the pulmonary arteries, veins and airways of PTL. The liver centerlines are our own, traced with VMTK from roots picked by hand; HiPaS and PARSE come as the centerline graphs of VesselTok [1], traced with the Voreen tool from their segmentation masks. The MouseLight axons come from NeuroMorpho.Org (RRID:SCR_002145) [2].

| Vessel | Dataset | Imaging | Centerlines | Cases | Specimens | Terminals | Branch points | Licence |
|---|---|---|---|---|---|---|---|---|
| Hepatic vein | VEELA [3] | portal-venous CTA of potential liver donors, 0.54–0.79 mm in plane, 2–3.2 mm slices | ours (VMTK) | 20 | 15 | 80 (45–105) | 1,078 | CC BY-NC-SA 4.0 (CHAOS images and challenge rules) |
| | LIRCAD [4] | CT of LiTS and 3Dircadb, resampled to isotropic 0.56–0.98 mm voxels | ours (VMTK) | 77 | 27 | 53 (41–104) | 1,476 | CC BY 4.0 |
| | HVA-CT [5] | CT of MSD Task 08, slices up to 2 mm | ours (VMTK) | 61 | 54 | 67.5 (45–87) | 3,341 | CC BY-SA 4.0 |
| Portal vein | VEELA | as above | ours (VMTK) | 20 | 17 | 64 (40–104) | 1,000 | as above |
| | LIRCAD | | ours (VMTK) | 77 | 24 | 54 (40–92) | 1,276 | |
| | HVA-CT | | ours (VMTK) | 61 | 34 | 56 (42–79) | 1,790 | |
| Pulmonary artery | PTL [6] | CT of 800 subjects from several centres, slices of 0.5–1.5 mm | release | 799 | 535 | 277 (200–460) | 147,172 | CC BY 4.0 |
| | HiPaS [7] | CT resampled to $0.65 \times 0.65 \times 1.0$ mm | VesselTok | 250 | 177 | 253 (200–387) | 43,496 | MIT; VesselTok CC BY-NC 4.0 |
| | PARSE [8] | CTPA of patients with lung nodules, 0.50–0.95 mm in plane, 1 mm slices | VesselTok | 100 | 92 | 310 (200–433) | 25,994 | by agreement; VesselTok CC BY-NC 4.0 |
| Pulmonary vein | PTL | as above | release | 799 | 646 | 295 (200–439) | 183,647 | as above |
| | HiPaS | | VesselTok | 250 | 198 | 264.5 (200–396) | 50,054 | |
| All vessels | | | | | 1,819 | | 460,324 | |
| Networks that are not blood vessels (Fig. 6 of the main text) | | | | | | | | |
| Human airway | PTL | as the PTL arteries | release | 799 | 799 | 55 (8–173) | 86,230 | CC BY 4.0 |
| Mouse airway | lapdMouse [9] | imaging cryomicrotome, 4.6 $\mu$m in plane, 9.52 $\mu$m slices | release | 34 mice | 34 | 373 (178–800) | 27,212 | CC BY 4.0 |
| Tomato shoot | Conn et al. [10, 11] | 3D laser scans, each plant on several days | release | 17 plants | 307 scans | 6 (2–35) | 1,796 | CC BY 4.0 |
| Arabidopsis shoot | Meng et al. [12], from Pan et al. [13] | point clouds carved from 72 photographs per scan | release | 5 plants | 52 scans | 17 (3–106) | 849 | CC BY 4.0 |
| Neuron axon | MouseLight [14] | two-photon block-face tomography, $0.3 \times 0.3 \times 1.0$ $\mu$m | release | 1,111 neurons | 1,112 | 135 (2–2,736) | 210,627 | CC BY 4.0 (NeuroMorpho.Org) |

#### References


[1] Prabhakar, C. *et al.* VesselTok: Tokenizing vessel-like 3D biomedical graph representations for reconstruction and generation. In *Computer Vision – ECCV 2026*, 131–149 (LNCS, 2026).

[2] Tecuatl, C., Ljungquist, B. & Ascoli, G. A. Accelerating the continuous community sharing of digital neuromorphology data. *FASEB BioAdvances* **6**, 207–221 (2024).

[3] Yazıcı, Z. A. *et al.* VEELA: A clinically-constrained benchmark for liver vessel segmentation in computed tomography angiography (2026). Preprint at `https://arxiv.org/abs/2605.22357`.

[4] Machta, Y. *et al.* Enhancing the automatic segmentation and analysis of 3D liver vasculature models (2025). Preprint at `https://arxiv.org/abs/2411.15778`.

[5] Salvi, M. *et al.* HVA-CT: Hepatic venous annotations CT. Zenodo, `https://doi.org/10.5281/zenodo.19850108` (2026).

[6] Xie, K., Yang, J., Wei, D., Weng, Z. & Fua, P. Efficient anatomical labeling of pulmonary tree structures via deep point-graph representation-based implicit fields. *Medical Image Analysis* **99**, 103367 (2025).

[7] Chu, Y. *et al.* Deep learning-driven pulmonary artery and vein segmentation reveals demography-associated vasculature anatomical differences. *Nature Communications* **16**, 2262 (2025).

[8] Luo, G. *et al.* Efficient automatic segmentation for multi-level pulmonary arteries: The PARSE challenge (2023). Preprint at `https://arxiv.org/abs/2304.03708`.

[9] Bauer, C., Krueger, M., Lamm, W. J. E., Glenny, R. W. & Beichel, R. R. lapdMouse: Associating lung anatomy with local particle deposition in mice. *Journal of Applied Physiology* **128**, 309–323 (2020).

[10] Conn, A., Pedmale, U. V., Chory, J. & Navlakha, S. High-resolution laser scanning reveals plant architectures that reflect universal network design principles. *Cell Systems* **5**, 53–62.e3 (2017).

[11] Navlakha, S. 3D scans of plant shoot architectures. Mendeley Data, `https://doi.org/10.17632/9k7zctdyhs.1` (2017).

[12] Meng, X., Piazza, B., Both, C., Barzel, B. & Barabási, A.-L. Surface optimization governs the local design of physical networks. *Nature* **649**, 315–322 (2026).

[13] Pan, H., Hétroy-Wheeler, F., Charlaix, J. & Colliaux, D. ARABIDOPSIS 3D+T dataset. Zenodo, `https://doi.org/10.5281/zenodo.5205562` (2021).

[14] Winnubst, J. *et al.* Reconstruction of 1,000 projection neurons reveals new cell types and organization of long-range connectivity in the mouse brain. *Cell* **179**, 268–281.e13 (2019).
